# CUX2 deficiency causes facilitation of excitatory synaptic transmission onto hippocampus and increased seizure susceptibility to kainate

**DOI:** 10.1101/2021.09.17.460803

**Authors:** Toshimitsu Suzuki, Tetsuya Tatsukawa, Genki Sudo, Caroline Delandre, Yun Jin Pai, Hiroyuki Miyamoto, Matthieu Raveau, Atsushi Shimohata, Iori Ohmori, Shin-ichiro Hamano, Kazuhiro Haginoya, Mitsugu Uematsu, Yukitoshi Takahashi, Masafumi Morimoto, Shinji Fujimoto, Hitoshi Osaka, Hirokazu Oguni, Makiko Osawa, Atsushi Ishii, Shinichi Hirose, Sunao Kaneko, Yushi Inoue, Adrian Walton Moore, Kazuhiro Yamakawa

## Abstract

*CUX2* gene encodes a transcription factor that controls neuronal proliferation, dendrite branching and synapse formation, locating at the epilepsy-associated chromosomal region 12q24 that we previously identified by a genome-wide association study (GWAS) in Japanese population. A *CUX2* recurrent *de novo* variant p.E590K has been described in patients with rare epileptic encephalopathies and the gene is a candidate for the locus, however the mutation may not be enough to generate the genome-wide significance in the GWAS and whether *CUX2* variants appear in other types of epilepsies and physiopathological mechanisms are remained to be investigated. Here in this study, we conducted targeted sequencings of *CUX2,* a paralog *CUX1* and its short isoform *CASP* harboring a unique C-terminus on 271 Japanese patients with a variety of epilepsies, and found that multiple *CUX2* missense variants, other than the p.E590K, and some *CASP* variants including a deletion, predominantly appeared in patients with temporal lobe epilepsy (TLE). The CUX2 variants showed abnormal localization in human cell culture analysis. While wild-type CUX2 enhances dendritic arborization in fly neurons, the effect was compromised by some of the variants. *Cux2*- and *Casp*-specific knockout mice both showed high susceptibility to kainate, increased excitatory cell number in the entorhinal cortex, and significant enhancement in glutamatergic synaptic transmission to the hippocampus. CASP and CUX2 proteins physiologically bound to each other and co-expressed in excitatory neurons in brain regions including the entorhinal cortex. These results suggest that *CUX2* and *CASP* variants contribute to the TLE pathology through a facilitation of excitatory synaptic transmission from entorhinal cortex to hippocampus.

## Introduction

*Cux2* gene encodes a homeobox transcription factor CUX2 that is predominantly expressed in progenitor cells of the subventricular zone in mouse embryos and pyramidal neurons of the upper neocortical layers (II–IV) in adult mice^1^. CUX2 is also expressed in Reelin-positive neurons distributed throughout the layers II–IV in postnatal day 0 (P0) mice^2^. CUX2 controls neuronal proliferation, dendrite branching, spine morphology and synapse formation^3,4^. We recently reported a genome-wide association study (GWAS) on 1,825 Japanese patients with variable epilepsies which identified an associated region at chromosome 12q24.11 – 12q24.13 harboring 24 transcripts including *CUX2* gene^5^. In the region, *CUX2* is only gene which has been reported to be relevant for epilepsy; a recurrent *de novo* variant (c.1768G>A, p.E590K) has been identified in patients with rare epileptic encephalopathies (EEs)^6,7^. *CUX2* is therefore one of the most promising candidate genes in this 12q24 epilepsy-associated region, but the mutation reported in rare EEs may not be enough to explain the association detected in the Japanese GWAS study.

To investigate whether *CUX2* and its paralogues' mutations are involved in other types of epilepsies, here we performed targeted sequencing of *CUX2*, its paralog *CUX1,* and *CASP* which is a short isoform of *CUX1* with a unique C-terminus, in Japanese patients with variable epilepsies including genetic generalized and structural/metabolic epilepsies, and identified their variants predominantly in patients with temporal lobe epilepsy (TLE), the most common but intractable form of epilepsy^8^. Analyses in human cultured cell and transgenic fly showed that the variants have loss-of-function effects. CUX2 and CASP deficiencies in mice increased their seizure susceptibilities to a convulsant, kainate, which has long been used to generate TLE animal models^9^. Histological and electrophysiological analyses revealed increases of excitatory neuron numbers in entorhinal cortex and those in excitatory input to hippocampus in both mice, proposing a circuit mechanism for the pathology of TLE.

## Materials and Methods

### Subjects

Genomic DNAs from 271 Japanese patients with a variety of epilepsies (Table S1) and 311 healthy Japanese volunteers recruited by cooperating hospitals were used for the targeted sequencing analyses for *CUX2*, CUX1 and *CASP*. For the frequency calculation of c.3847G>A (p.E1283K) variant in *CUX2* gene, additional DNA samples from independent 69 Japanese patients with TLE from 2 additional independent facilities were further recruited (Table S1). The patients' DNAs analyzed in our GWAS^5^ were not used in the present study, because their epilepsy subtype information [TLE, etc.] were not available for those materials.

### Targeted sequencing

Genomic DNAs were extracted from peripheral venous blood samples using QIAamp DNA Blood Midi Kit (Qiagen). Genomic DNA samples were amplified with the illustra GenomiPhi V2 DNA Amplification Kit (GE Healthcare). We designed PCR primers to amplify all coding regions of *CUX2* (NM_015267), *CUX1* (NM_001202543 and first exon of NM_181552), unique C-terminus region of *CASP* (NM_001913), and amplified genomic DNA by PCR using the PrimeSTAR HS DNA Polymerase (TaKaRa) or KOD-plus Ver. 2 (TOYOBO). The PCR products were purified using ExoSAP-IT PCR product Cleanup (Thermo Fisher Scientific) and analyzed by direct sequencing using the ABI PRISM 3730xl Genetic Analyzer. All novel variants identified in amplified DNA by GenomiPhi were verified by direct-sequencing of patients’ genomic DNAs. Primer sequences and PCR conditions are available upon request.

**Quantification of mRNA –** described in the Supplemental Methods.

### Domain search

Domain searches in CUX2, CUX1, and CASP amino acid sequences were performed using the SMART database.

**Expression constructs and mutagenesis –** described in the Supplemental Methods.

**Cell imaging –** described in the Supplemental Methods.

**Drosophila stocks and crosses –** described in the Supplemental Methods.

**TUNEL assay in flies –** described in the Supplemental Methods.

### Mice

*Cux2* knockout (KO) mouse was obtained from Texas A&M Institute for Genomic Medicine (TIGM) as cryopreserved sperm of heterozygous *Cux2* gene trap mouse (129/SvEv × C57BL/6 background) derived from the gene trapped clone OST440231. Live animals were produced by *in vitro* fertilization at the Research Resources Division (RRD) of the Institute of Physical and Chemical Research (RIKEN) Center for Brain Science. The heterozygous mice were maintained on the C57BL/6J (B6J) background, and the resultant heterozygous mice were interbred to obtain wild-type (WT), heterozygous, and homozygous mice. Genotyping was carried out as described previously^10^.

*Casp*-specific KO mice were generated using the CRISPR/Cas system. Plasmid vector pX330-U6-Chimeric_BB-CBh-hSpCas9 was a gift from Dr. Feng Zhang (Addgene plasmid # 42230). A pair of oligo DNAs corresponding to *Casp*-gRNA (TTTCCATCATCTCCAGCCAA AGG) in exon 17 of *Casp* (NM_198602) was annealed and ligated into pX330-U6-Chimeric_BB-CBh-hSpCas9. For knock-out mouse production, Cas9 mRNAs and *Casp*-gRNA were diluted to 10 ng/μL each. Further, B6J female mice and ICR mouse strains were used as embryo donors and foster mothers, respectively. Genomic DNA from founder mice was extracted, and PCR was performed using gene-specific primers (CRISPR_check_F: 5’-GGAGCTATTGTAGGACATCACAGA-3’ and CRISPR_check_R: 5’-CCCCAGTGTTCTTTACTTTGAGTT-3’). PCR products were purified using ExoSAP-IT PCR product Cleanup and analyzed by direct sequencing using the ABI PRISM 3730xl Genetic Analyzer. The heterozygous mutant mice (c.1514^1515ins.TT, p.S506fs) were crossbred with B6J mice, and the resultant heterozygous mice were interbred to obtain WT, heterozygous, and homozygous mutant mice. The frame-shift mutation was confirmed by sequence analysis of cDNA from mutant mouse brains.

**Seizure susceptibility in mice –** described in the Supplemental Methods.

### CUX2 antibody generation

To generate a rabbit polyclonal antibody to CUX2, a fusion protein was prepared, in which the glutathione-S-transferase (GST) protein was fused to the a.a. residues 356 to 415 of mouse CUX2 which has been used in the previous study's antibody generation^11^. Rabbits were injected with 0.5 mg of purified GST fusion protein in Freund’s complete adjuvant, boosted five times with 0.25 mg of protein, and serum collection at 1 week following the last boost. Polyclonal antibody was purified by affinity chromatography. The serum was passed through a GST affinity column ten times, and the flow-through was then applied to a GST-CUX2 (356-415 a.a.) affinity column to isolate antibodies.

**Histological analyses –** described in the Supplemental Methods.

**In vitro electrophysiology –** described in the Supplemental Methods.

**Co-immunoprecipitation –** described in the Supplemental Methods.

### Statistical analysis

In the *in vitro* and *in vivo* experiments, data are presented as Box-and-whisker plots or means ± s.e.m. The boxes show median, 25th and 75th percentiles, and whiskers represent minimum and maximum values. *P*-value for p.E1283K was calculated using the Cochrane-Armitage trend test. One-way or two-way ANOVA, Tukey’s multiple comparison test, Chi-square test, or Kolmogorov-Smirnov test were used to assess the data as mentioned in the figure legends.

Statistical significance was defined as *P* < 0.05.

### Study approval

#### Human study

The experimental protocols were approved by the Ethical Committee of RIKEN and by the participating hospitals and universities. All human study experimental procedures were performed in accordance with the guidelines of the Ethical Committee of the RIKEN and with the Declaration of Helsinki. Written informed consents were obtained from all individuals and/or their families in compliance with the relevant Japanese regulations.

#### Animal study

All animal experimental protocols were approved by the Animal Experiment Committee of RIKEN. All animal breeding and experimental procedures were performed in accordance with the ARRIVE guidelines and the guidelines of the Animal Experiment Committee of the RIKEN.

### Data availability

All data generated or analyzed during this study are included in this published article and its Supplementary Information File.

## Results

### *CUX2* variants predominantly appeared in Japanese TLE patients

We carried out a targeted sequencing of *CUX2* in 271 Japanese patients with variable epilepsies consisting of 116 genetic generalized epilepsies and 155 structural/metabolic ones (Table S1). Structural/metabolic epilepsy samples contained 68 TLEs, which were further divided to 57 mesial TLE (mTLE) and 11 lateral TLE (lTLE). We identified five *CUX2* heterozygous missense variants in nine unrelated patients (Figure 1A, Table 1 and Supplemental Note). Notably, eight of the nine patients with *CUX2* variants had TLE (one lTLE and seven mTLE patients). All of patients carrying *CUX2* missense variants belonged to the subgroup of structural/metabolic epilepsy. None of patients with genetic generalized epilepsies showed *CUX2* variants except for silent variants. All of the mTLE patients showed hippocampal sclerosis. Three (p.R34W, p.P454L, and p.W958R) out of the five variants were absent or rare (< 0.5%) in the in-house Japanese control individuals (in-house controls) and databases and were also predicted to be damaging (Table 1). The p.E1283K variant, a frequent variant predicted to be less damaging, appeared in Japanese TLE patients at a significantly high ratio [*P* = 5.93 × 10^−3^, OR = 6.94, 95% CI = 1.39–34.61 calculated in 137 (above-mentioned 68 + additional independent 69; Table S1) TLE patients vs 311 in-house controls] and therefore we hypothesized it may also be a genetic contributor for TLE. The p.D337N variant appeared in one case with TLE and controls with a similar allele frequency.

**Figure 1.**
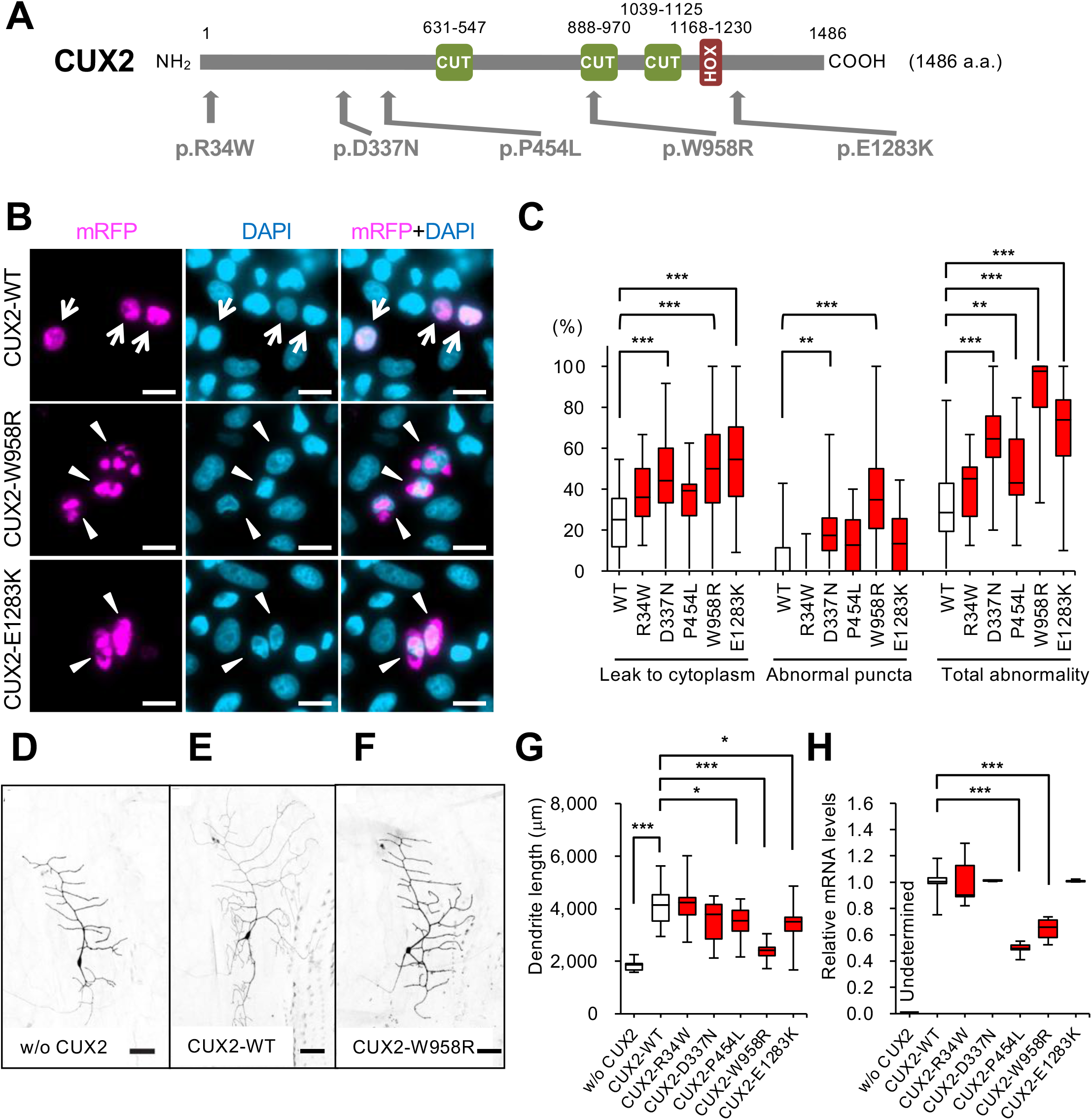
Loss-of-function effects of TLE variants in *CUX2*. **(A)** CUX2 protein structure (NP_056082) with variants appeared in patients with epilepsy. (**B**) Abnormal subcellular localization of CUX2 variant proteins. CUX2-WT protein (arrows) was limited to, but well distributed within, nuclei stained with DAPI (cyan), whereas variants showed abnormal aggregates in nuclei (W958R) or leaked-out to the cytoplasm (E1283K) (arrowheads). Scale bars = 20 μm. (**C**) Ratio of abnormally localized CUX2 proteins (> 200 cells counted). n = WT: 545, R34W: 282, D337N: 363, P454L: 239, W958R: 565, and E1283K: 323 cells. (**D-H**) CUX2 WT accelerated arborization of fly neurons and TLE variants lowered its activity and expression. Representative images of neurons without CUX2 (D), WT control (E), and W958R (F). Scale bars = 50 μm. (G) Shortened dendrite length in transgenic fly with mutants (n = 11–25 neurons per genotypes) and (H) lowered expression of mutants (n = 6). One-way ANOVA Tukey’s multiple comparison test (C, G, H). *P < 0.05, **P < 0.01, ***P < 0.001.

**Table 1:**
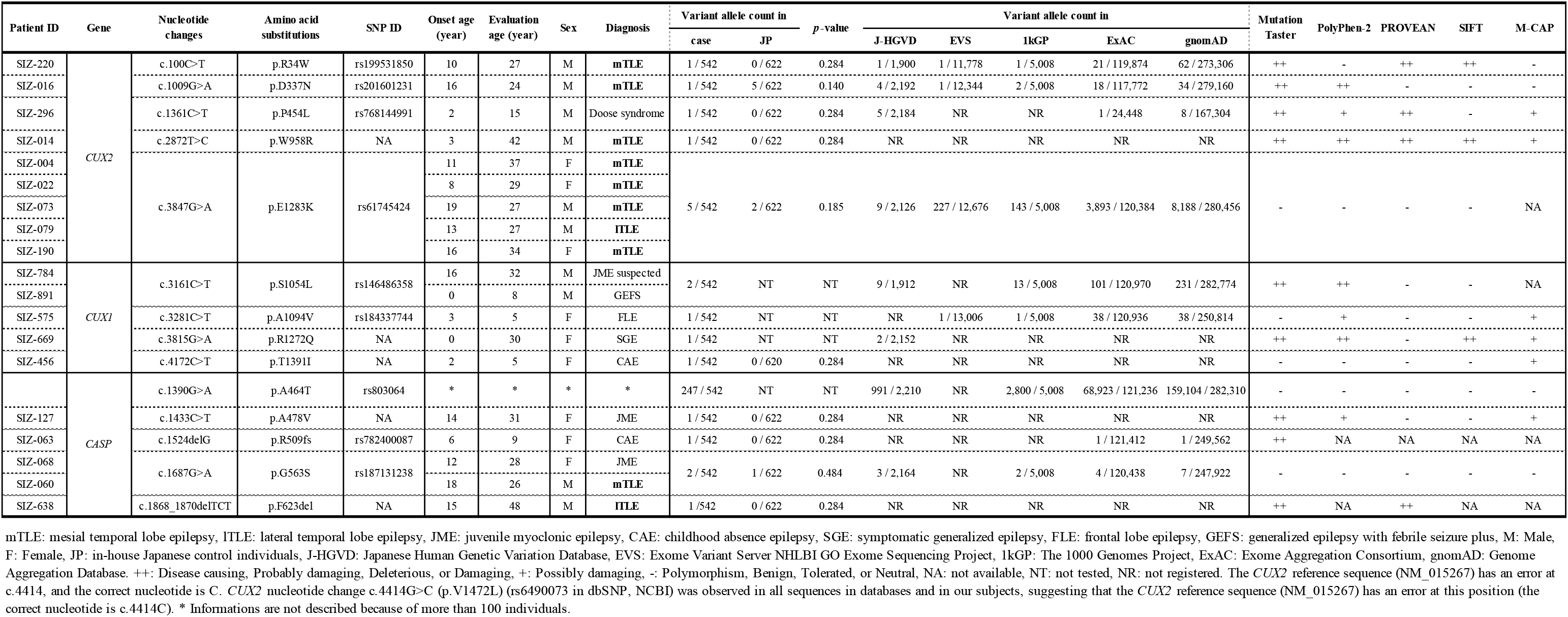
*CUX2*, *CUX1*, and *CASP* gene nonsynonymous variants in patients with epilepsy.

### Loss-of-function effects of *CUX2* variants

To investigate the functional impacts of *CUX2* variants appeared in patients with epilepsy (Figure 1A, Table 1), we transfected HeLa.S3 cells with expression constructs of wild-type (WT) and the five variants (p.R34W, p.D337N, p.P454L, p.W958R, and p.E1283K). We calculated two parameters of abnormality, “leakage to cytoplasm” and “abnormal puncta” (Figure 1B, C and Figure S1A). Although some but not all variants showed abnormalities in each parameter, the combined data reached statistical significance except for p.R34W.

*CUX2* is an ortholog of *Drosophila melanogaster cut,* which promotes dendritic arbor morphological complexity^12^. We generated transgenic fly lines to express human WT CUX2 or variants (p.R34W, p.D337N, p.P454L, p.W958R, and p.E1283K) and analyzed their dendritic arbor morphology in *Drosophila* larvae (Figure 1D-G and Figure S1B-D). Similar to its *Drosophila* orthologue^12^, ectopic expression of CUX2 WT strongly increased dendritic arbor complexity (branch number and length). However, activities to drive arbor complexity in the variants were significantly decreased, except for p.R34W and p.D337N. RT-qPCR assays in the adult transgenic flies revealed that expression levels of the *CUX2* variants, p.P454L and p.W958R, were significantly lower (Figure 1H). All *CUX2* constructs were inserted into the same genomic site, therefore the lower expression levels of transgenes are not likely to be due to position effects but most likely due to these variants because these are only the differences in the constructs used for the analyses of fly. TUNEL of adult fly brains showed that the alleles did not promote apoptotic cell death (Figure S1E). Together, these observations suggest that the *CUX2* variants present in patients with epilepsy cause loss-of-function of the protein.

### *Cux2*-deficient mice show increased susceptibility to kainate

Because of the loss-of-function nature of epilepsy-associated *CUX2* variants, we next investigated *Cux2-*KO mice^10^. The body weight of 2-month-old mice was comparable among genotypes (Figure S2A). In electrocorticogram analysis, the median of the poly spike and wave discharges frequency was slightly higher in the primary somatosensory cortex forelimb region of *Cux2*(−/−) than WT mice, however the difference did not reach statistical significance (data not shown). No obvious epileptic behaviors or changes in local field potential recordings in the hippocampus were noted in *Cux2*(+/−) or *Cux2*(−/−) mice. Although patients with TLE often have past histories of febrile seizures^13^, *Cux2*-KO mice did not show any seizure susceptibility to increased body temperature (data not shown). Seizure susceptibility to pentylenetetrazole (PTZ), a GABA-A receptor antagonist, remained unchanged in *Cux2*-KO mice (Figure S2B-F). Importantly, however, *Cux2*-KO mice had a high susceptibility to kainate, which is commonly used to generate TLE animal models^9^, in frequencies of generalized convulsive seizures (GS) (Figure 2A) and lethality (Figure 2B). The latencies to onset of GS and death were also significantly decreased in *Cux2*(−/−) mice (Figure S2G, H). Seizure severity was also significantly higher in *Cux2*-KO female mice (Figure 2C). These results support the notion that *CUX2* loss-of-function mutations cause TLE.

**Figure 2.**
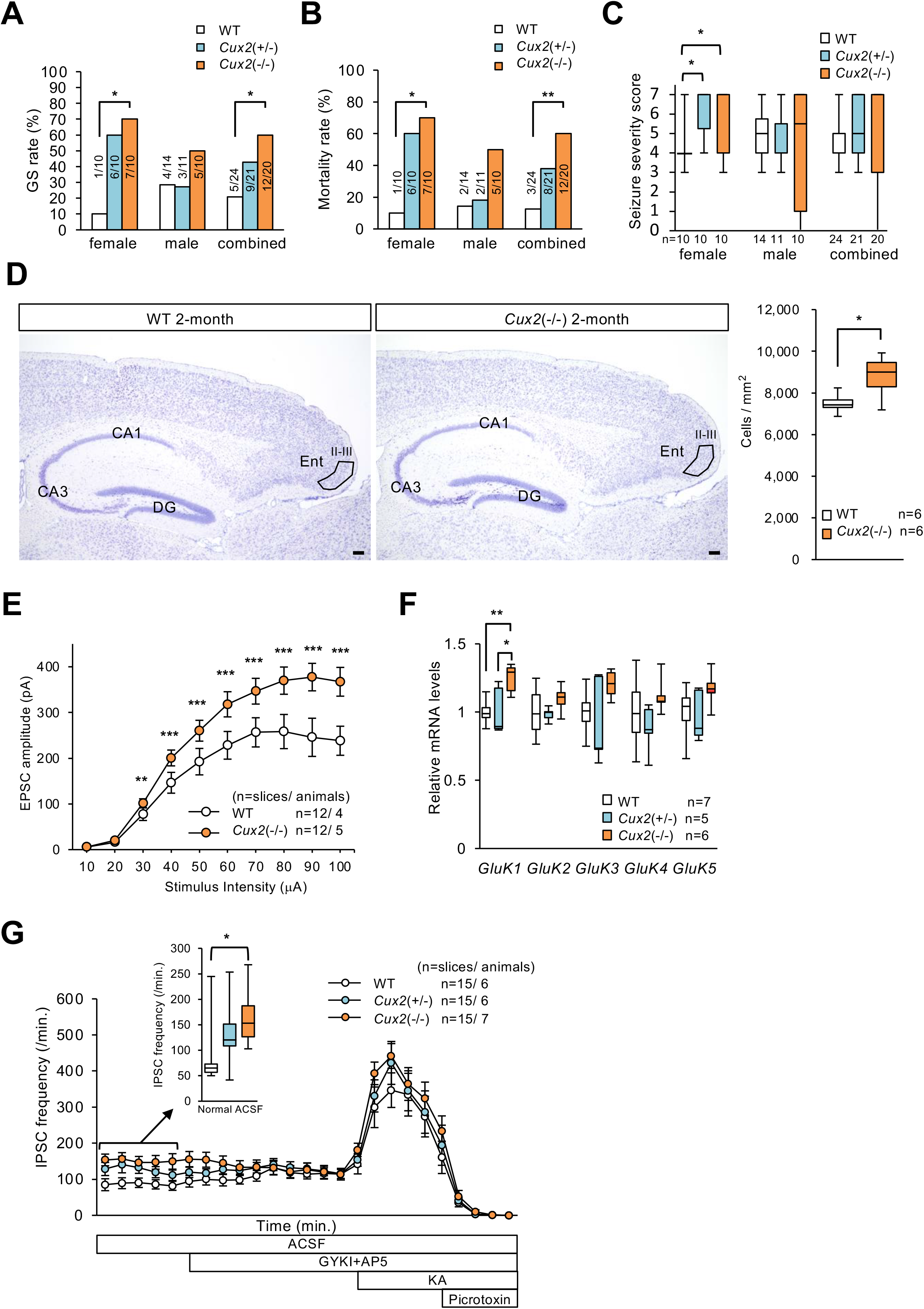
Increased kainate susceptibility, entorhinal cortical cell number, and excitatory input to hippocampal granule cells in *Cux2*-KO mice. **(A-C)** Seizure-related events in mice after intraperitoneal injection of kainate (KA). Ratio of animals exhibiting generalized convulsive seizure (GS) (A), mortality rate (B), and seizure severity scores (C) was significantly higher in *Cux2*(−/−) female and combined gender mice. **(D)** Number of entorhinal cortex layer II-III excitatory neurons was significantly increased in *Cux2*(−/−) mice (2-month-old). Scale bar = 100 μm. **(E)** Slice-patch analyses showed that perforant path-evoked EPSCs in dentate granule cells were significantly increased in *Cux2*(−/−) female. **(F)** RT-qPCR analyses revealed that *GluK1* mRNA was significantly increased in *Cux2*(−/−) mice. **(G)** Basal frequency of sIPSC in dentate granule cells of *Cux2*(−/−) female was significantly increased, and it was suppressed with subsequent applications of antagonists for AMPA receptor (GYKI) and NMDA receptor (AP5). Kainate (KA) increased the sIPSC frequency, which was then suppressed by the GABA-A receptor antagonist picrtoxin. DG; dentate gyrus, Ent; entorhinal area. Yates' correction after Pearson’s Chi-square (A, B), one-way ANOVA Tukey's test (C, F, G), one-way ANOVA (D), or two-way ANOVA Tukey's test (E). n: mouse numbers. *P < 0.05, **P < 0.01, ***P < 0.001.

### *Cux2*-deficient mice show increased cell number in entorhinal cortex and glutamatergic input to hippocampus

*Cux2*(−/−) mice have been reported to show overgrowth of the neocortical upper layers^3^. In a Nissl staining, we also found a significantly increased cell number in entorhinal cortex layers II-III, which projects to hippocampal dentate granule cells and CA3 pyramidal cells (Figure 2D). In the slice-patch recordings, we further found that perforant path-evoked excitatory postsynaptic currents (eEPSCs) in the dentate granule cells were significantly higher in *Cux2*(−/−) mice (Figure 2E), indicating that glutamatergic synaptic transmission from the entorhinal cortex layers II–III onto the hippocampus was significantly facilitated in *Cux2*-KO mice.

At a glance, hippocampal structures in *Cux2*-KO mice were comparable to 2- and 10-month-old WT mice (Figure 2D, Figure S3). We generated an anti-CUX2 antibody similarly to a previous study^11^ and confirmed the presence of CUX2 immunosignals in WT and absence in *Cux2*(−/−) mice (Figure S3A, B). CUX2 immunosignals were dense in the neocortical upper layers (II-IV) as previously reported^1^ and also dense in the entorhinal cortex upper layers (II-III) (Figure S3C). In WT hippocampus, we only observed CUX2 immunolabeling signals in inhibitory interneurons, specifically somatostatin (SST)-positive, reelin (RLN)-positive and parvalbumin (PV)-positive inhibitory, but not in excitatory neurons (Table S2, Figure S4). We found that there were no significant differences in interneuron cell numbers between genotypes (Figure S5A-D). Timm staining and immunohistochemistry for c-Fos (Figure S5E, F), Doublecortin, phospho-Histone H3, Ki67, NeuN, GFAP, and ZnT-3 (data not shown) did not show differences in the hippocampus between genotypes. There are five subtypes of kainate receptors (KARs), GLUK1–GLUK5, in primates and rodents. We investigated KARs expression in the hippocampus of 2-month-old *Cux2*-KO female mice. RT-qPCR assays revealed that the expression of *GluK1* (formerly named *GluR5*) was significantly higher in *Cux2*(−/−) mice (Figure 2F), which is presumably a homeostatic compensatory reaction to epileptic seizures (see Discussion). The baseline frequencies of spontaneous inhibitory postsynaptic currents (sIPSCs) in hippocampal dentate granule cells were significantly higher at 6–7 weeks old, and this difference was suppressed after bath-application of GYKI and AP5, which are AMPA and NMDA receptor antagonists, respectively (Figure 2G). Frequency of sIPSC in dentate PV-positive interneurons was also increased in *Cux2*-KO mice (Figure S6). These results suggest that, in *Cux2*(−/−) mice, the function of hippocampal inhibitory neurons remained intact, but the increased excitatory input from the entorhinal cortex to the hippocampus could facilitate firing activities of inhibitory neurons, which itself would also be a compensatory action to epileptic activities in mice. In CA3 pyramidal neurons of *Cux2*-KO mice, EPSCs were not significantly affected (Figure S6), suggesting that the increased excitatory input in the upstream dentate granule cells may be neutralized by the increased inhibitory input in those cells.

Taken together, these results suggest that the increase in entorhinal cortex layers II–III cell numbers and the resultant facilitation of glutamatergic synaptic transmission from the entorhinal cortex layers II–III onto hippocampi are causal factors leading to the increased susceptibility to kainate of *Cux2*-KO mice. Increases in GLUK1 and facilitated firing of inhibitory neurons in the mouse hippocampus would rather be compensatory reactions.

### *CASP* variants in TLE patients

*CUX1* is a paralog of *CUX2*, and *CASP* is an alternatively spliced short isoform of *CUX1* harboring a unique C-terminus^14^ (Figure 3A, Figure S7). CUX1 and CUX2 proteins have four DNA binding domains (three CUT repeats and one homeodomain), but CASP lacks all of these domains and instead contains a transmembrane domain. We performed targeted sequencing analyses of *CUX1* and *CASP* in the 271 Japanese patients with epilepsy and identified nine nonsynonymous variants (Figure 3A, Table 1, and Supplemental Note). Among those, one variant in *CUX1*, c.4172C>T (p.T1391I) and three variants in *CASP*, c.1433C>T (p.A478V), c.1524delG (p.R509fs), and c.1868_1870delTCT (p.F623del), were completely absent or very rare in in-house controls and databases (Table 1). No other truncation variants of the *CASP*-specific sequence were found in these databases, suggesting that *CUX1* and especially *CASP* variants contribute to epilepsy. Although epilepsies observed in patients with *CUX1* and *CASP* variants were rather heterogeneous, CASP-p.G563S and p.F623del variants appeared in mTLE and lTLE patients, respectively (Table 1). The mTLE patient SIZ-060 showed hippocampal sclerosis (Supplemental Note).

**Figure 3.**
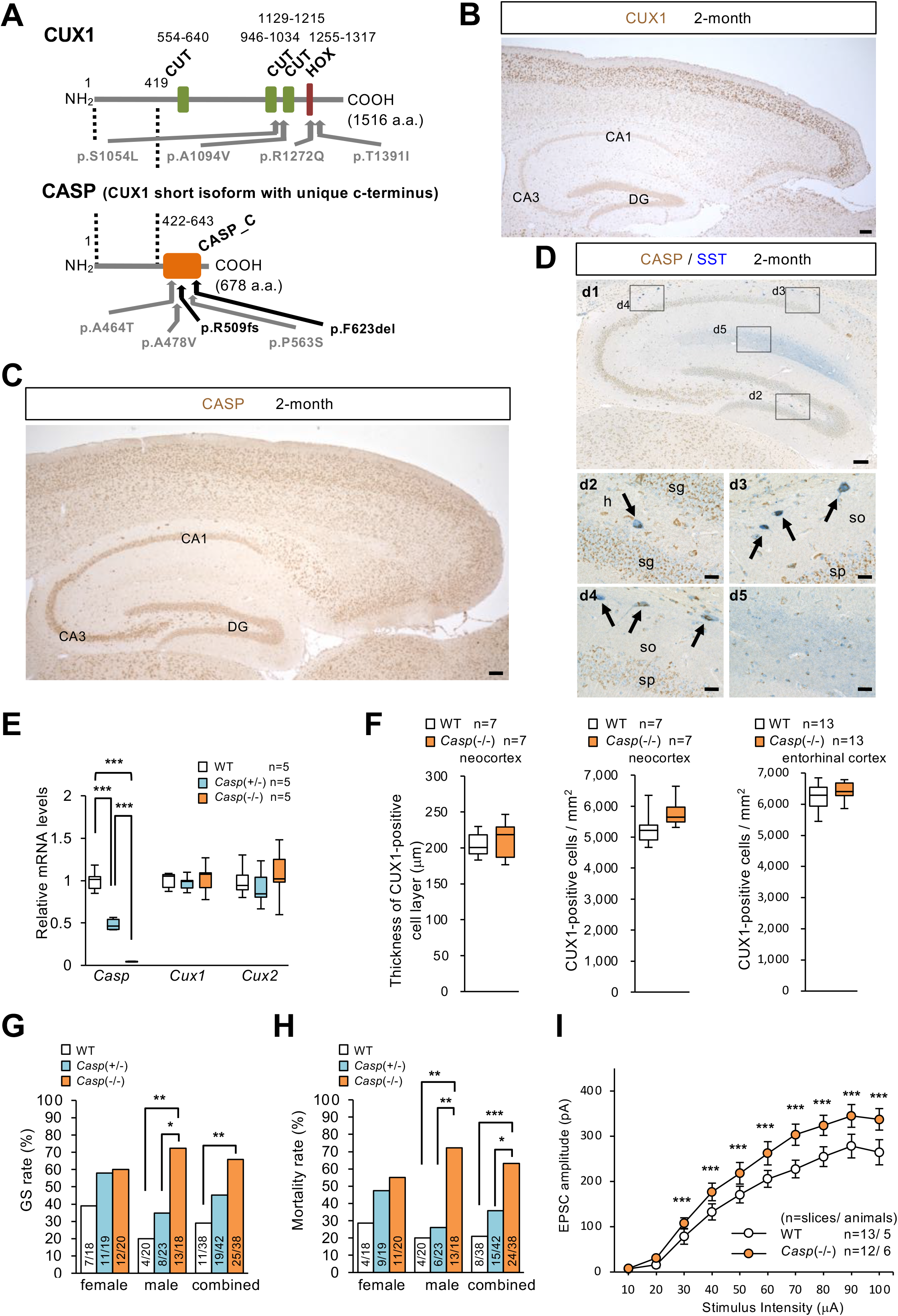
*CASP* variants in epileptic patients, CASP distribution, and increases in kainate susceptibility and excitatory input to hippocampal granule cells in *Casp*-KO mice. **(A)** Locations of *CUX1* and *CASP* variants in patients with epilepsy (see Table 1). Dashed lines define the common region. **(B)** CUX1 immunosignals (brown) in neocortical and entorhinal cortex upper layer excitatory neurons and hippocampal interneurons. **(C)** CASP (brown) expressed more widely in neurons, and intensely expressed in neocortical and entorhinal cortex upper layer excitatory neurons. **(D)** In hippocampus, CASP (brown) was dense in SST-positive (blue) interneurons at hilus and stratum oriens (arrows). d2-d5; magnified images outlined in d1. Scale bars = 100 μm (B, C and d1), 20 μm (d2-d5). so; stratum oriens, sp; stratum pyramidale, sg; stratum granulosum, h; hilus. **(E)** RT-qPCR analyses revealed that *Casp* mRNA was decreased, while *Cux1* and *Cux2* mRNAs remained unchanged, in *Casp*-KO mice. **(F)** Thickness of the CUX1-positive neocortical layer (left), density of CUX1-positive cell in neocortex (middle), and CUX1-positive cell density in entorhinal cortex (right). CUX1-positive cell density tended increase at neocortex and entorhinal cortex in *Casp*(−/−) mice (2-month-old) but not statistically significant. **(G, H)** *Casp*-KO mice showed significantly higher susceptibility to kainate in seizure rate (G), mortality (H). **(I)** Perforant path-evoked EPSCs in dentate granule cells were significantly increased in *Casp*(−/−) male (6–7-week-old). One-way ANOVA Tukey's test (E), Yates' correction after Pearson’s Chi-square (G, H), or two-way ANOVA Tukey's test (I). n: mouse numbers. *P < 0.05, **P < 0.01, ***P < 0.001.

### CASP and CUX2 proteins are co-expressed in excitatory neurons of entorhinal cortex upper layer and physiologically bind to each other

Immunohistochemistry with CUX1 antibodies in 2-month-old WT mice revealed CUX1 immunosignals in excitatory neurons at the neocortical upper layers (II-IV), as previously reported^3^, and those at the entorhinal cortex upper layers (II-III) (Figure 3B), similar to CUX2 (Figure S3C). In contrast in the hippocampus, CUX1 was expressed in SST-positive, RLN-positive, and PV-positive interneurons, but not in excitatory neurons (Figure 3B and Figure S8), similar to CUX2 (Figure S4). Using a CASP-specific antibody recognizing 400–650 a.a., we found that CASP was rather widely expressed in neurons of multiple brain regions, but still dense in the neocortical and entorhinal cortex upper layers, similar to CUX1 and CUX2 (Figure 3C). In the hippocampus, CASP was dense in hilar and stratum-oriens SST-positive cells that expressed CUX2 (Figure 3C, D), and more specifically, within the cytoplasm (Figure S9), which is consistent with CASP expression in the Golgi apparatus^15^.

A protein interaction between CUX1 and CASP has been previously reported^14^. Here we newly found that the CASP protein physically interacts with CUX2 (Figures S10 and S11). All three tested CUX2 rare variants bound to CASP, and all three tested CASP rare variants bound to CUX2 (Figures S10 and S11), suggesting that the variants did not affect protein binding between CASP and CUX2.

### *Casp*-deficient mice also show increased cell number in entorhinal cortex and glutamatergic input to hippocampus

It has been reported that the number of cortical neurons was significantly increased in *Cux1*(−/−); *Cux2*(−/−) double-mutant mice, but this increase was no greater than that in the *Cux2*(−/−) single mutant; therefore, regulation of the upper layer neuronal number was assumed to be a unique function of CUX2 and not redundant with CUX1 activities^3^. Because of the low survivability of *Cux1*-KO mice^16^ and our observation that the TLE variants appeared in the *CASP*-unique sequence but not in *CUX1* itself, we decided to investigate *Casp*-specific KO rather than *Cux1*-KO mice for analysis. We generated a *Casp-*specific KO mouse by targeting exon 17 at the unique C-terminus (Figure S7C). *Casp*(+/−) and *Casp*(−/−) mutant mice were born at a Mendelian ratio, grew normally, and were fertile. RT-qPCR analyses revealed that the *Casp* mRNA became half and diminished in *Casp*(+/−) and *Casp*(−/−) mice, respectively, whereas *Cux1* and *Cux2* mRNA levels remained unchanged (Figure 3E). CASP immunosignals well disappeared in *Casp*(−/−) mice (Figure S9A), confirming the specificity of the CASP antibody. At a glance, there were no abnormal localizations and intensities of CUX1 and CUX2 proteins in *Casp*-KO mice (Figure S9B, C). The median body weight was comparable among genotypes at 2 months of age (Figure S9D). RT-qPCR assays of KARs mRNA in the hippocampi of 2-month-old *Casp*-KO mice did not show significant change in KARs expression levels (Figure S9E).

In a Nissl staining of 2-month-old *Casp*-KO mice, no increase of neuron number was observed in the entorhinal cortex (Figure S9F). However, immunohistochemical staining using the anti-CUX1 antibody as a marker of neurons at upper layers of the neocortical (II–IV) and the entorhinal cortex (II–III) showed a tendency of increase in both the neocortex and entorhinal cortex (*P* = 7.57 × 10^−2^ and *P* = 4.50 × 10^−1^, respectively) (Figure3F). Furthermore, *Casp*-KO mice also showed high susceptibility to kainate (Figure 3G, H, Figure S9G-I). After intraperitoneal application of kainate, a larger number of *Casp*-KO mice showed GS and lethality (Figure 3G, H). Onset latencies of GS and death were significantly decreased in *Casp*-KO mice (Figure S9H, I). Seizure severity in *Casp*(−/−) mice was also significantly higher (Figure S9G). The differences in seizure susceptibility to kainate were seen mainly in male *Casp*-KO mice (Figure S9), contrary to *Cux2*-KO mice in which the susceptibility is higher in female (Figure S2). Notably again, perforant path-evoked EPSCs (eEPSCs) in the dentate granule cells were significantly higher in *Casp*-KO mice (Figure 3I), which is similar to *Cux2*-KO mice (Figure 2E).

All of these observations propose that facilitation of glutamatergic synaptic transmission from the entorhinal cortex onto hippocampal dentate granule cells is a common mechanism for TLE caused by *CUX2* and *CASP* variants.

## Discussion

In this study, we performed targeted sequencing analyses of *CUX2, CUX1* and *CASP* on 271 Japanese patients with a variety of epilepsies, and found that *CUX2* missense variants predominantly appear in TLE patients, in that eight of 68 TLE patients (12%) had *CUX2* variants. Three variants (p.R34W, p.P454L, and p.W958R) are quite rare or even absent in various databases, and are consistently predicted to have a damaging effect, therefore these would be regarded as causal or large-effect susceptibility variants. Although the p.E1283K is a high-frequent, relatively common variant, its frequency in TLE patients is statistically higher compared with in-house controls and therefore would potentially be a genetic contributor for TLE. *CASP* variants also appeared in two TLE patients. All of these patients with *CUX* family variants showed symptoms of epileptic seizures, suggesting that the variants may contribute to the threshold for the triggering epileptic seizures through a facilitation of excitatory synaptic transmission from entorhinal cortex to hippocampus in epilepsies caused by *CUX* family variants. Our recent GWAS analysis of Japanese patients with variable epilepsies identified a region with genome-wide significance at chromosome 12q24 which harbors *CUX2*^5^. Although a recurrent *de novo CUX2* variant p.E590K has been described in patients with EEs^6,7^, a previous whole exome sequencing study for Japanese patients with EEs^17^ did not find the *CUX2* pathogenic variant. Therefore, the *CUX2* recurrent variant in EEs may not be enough to explain the genome-wide association with epilepsy at the 12q24 region in Japanese population. In our GWAS study^5^, sub-analyses for subtypes of epilepsies further revealed that a polymorphic marker at the 12q24 epilepsy-associated region showed genome-wide significant association with structural/metabolic epilepsy. Hippocampal sclerosis is the major entity for structural/metabolic epilepsy, and therefore the *CUX2* variants in patients with TLE would contribute to the association with epilepsy at 12q24 in Japanese population.

Human cell culture and fly dendritic arborization analyses revealed loss-of-function effects of the *CUX2* variants, which were found in TLE patients. *CASP* also showed variants in epilepsy patients including TLE at the unique C-terminus and we further found that the CASP physically binds to CUX2. Although all tested CASP variants did not affect the binding activity to CUX2, the CASP protein has been reported to play a role in intra-Golgi retrograde transport^18^ and therefore the variants in CASP may still affect the subcellular transport or protein modification of CUX2.

*Cux2*- and *Casp*-KO mice did not show spontaneous seizures but showed significantly elevated seizure susceptibility to kainate, an agent which has been used to establish TLE animal models^9^. We previously reported a nonsense mutation of the *KCND2* gene encoding a voltage-gated potassium channel Kv4.2 in a patient with TLE^19^. Similar to *Cux2*- and *Casp*-KO mice, *Kcnd2*-KO mice did not show spontaneous epileptic seizures but showed increased susceptibility to kainate in seizure and mortality^20^. These observations suggest that increased seizure susceptibility to kainate correlates with the threshold for triggering epileptic seizures. However, other additional as-yet unknown modifying, genetic, or environmental factors may influence full expression of TLE symptoms including hippocampal sclerosis.

*Cux2*-KO mice showed a significant and *Casp*-KO mice showed a tendency of, increases in entorhinal cortex layer II–III excitatory neuron cell number. Although *Casp*-KO mice did not show any significant changes in *Cux2* mRNA expression levels and histological or cytological distributions of CUX2 protein, co-expression of CASP and CUX2 proteins in neurons including entorhinal cortex projection neurons, and the physiological interaction between CASP and CUX2 proteins still suggest that CASP deficiency may impair CUX2 function through an as-yet unknown mechanism, consequently leading to increased entorhinal cortex excitatory neuron cell number in *Casp*-KO mice. Furthermore, both *Cux2*- and *Casp*-KO mice revealed significant increases in perforant path-evoked EPSCs in dentate granule cells. These results suggest that facilitation of glutamatergic synaptic transmission from the entorhinal cortex onto hippocampal dentate granule cells is a causal basis for the significant increase in seizure susceptibilities to kainate in *Cux2*- and *Casp*-KO mice. In contrast, the observed changes in the hippocampus of *Cux2*-KO mice are assumed to be homeostatic compensatory reactions to the epileptic causal changes. In the hippocampus of WT mice, CUX2 immunolabelling was only observed in inhibitory interneurons such as SST-positive, RLN-positive, and PV-positive inhibitory, but not in excitatory neurons. In the *Cux2*-KO mice, although no changes were observed in interneuron cell numbers, a significant increase in sIPSC frequency was observed in dentate granule cells similar to patients with TLE^21^, which would presumably be a compensatory reaction^22,23,24^ to the increased excitatory input from the entorhinal cortex to the hippocampus. The increased GLUK1 expression in the hippocampus of *Cux2*-KO mice may also be suppressive for epileptic seizures because *GluK1* is expressed in inhibitory neurons^25^, and GLUK1 expression has been assumed to be protective at least for kainate-induced epilepsy^26^. Taken together, the changes within the hippocampus of *Cux2*-KO mice may be homeostatic compensatory responses rather than causal actions to epileptic seizures, and these changes in the hippocampus themselves also support the occurrence of epileptic causal changes in these mice.

In summary, our results of mutation analyses of *CUX* family genes in patients with epilepsies including TLE and the functional and mouse model analyses suggest that *CUX* family gene deficiency is one of the bases for TLE and that increase of cell number in the entorhinal cortex projection neurons and resultant increase of glutamatergic synaptic transmission to hippocampus is a possible pathological mechanism for TLE. Further investigations using mouse models with heterozygous missense variants, which have been identified in TLE patients are required to clarify whether the variants are true loss-of-function mutations and contribute to the TLE pathology.

## Acknowledgments

We thank the patients, families, volunteers, Lab for Genetic Control of Neuronal Architecture, Lab for Neurogenetics, Research Resources Division, Center for Brain Science, RIKEN. The authors also thank Dr. A. Miyawaki for the mRFP plasmid. This work was partly supported by grants from the Center for Brain Science and President's Discretionary Fund, RIKEN, and the Japan Agency for Medical Research and Development (AMED) under Grant Number JP20ek0109388.

## Author contributions

TS and KY designed the experiments; TS, TT, GS, CD, YJP and MR performed statistical analyses; TS, IO, SH, KH, MU, YT, MM, SF, H. Osaka, H. Oguni, MO, AI, SH, SK, YI and KY managed DNA samples; TS, IO, SH, KH, MU, YT, MM, SF, H. Osaka, H. Oguni, MO, AI, SH, SK, YI and KY recruited case and control samples; TS and KY performed targeted sequencing analyses; TS, CD, YJP, AWM, and KY performed functional analyses; TS, GS, TT, HM, AS, and KY performed mouse analyses; and TS, RM and KY wrote the manuscript.

## Competing Interests

The authors declare no competing interests.

## Supplementary Figure Legends

**Figure S1. CUX2 mutants show abnormal subcellular localizations in human. cultured cells and decreased branching effects but no apoptosis in fly.** (**A**) In HeLa.S3 cells, CUX2 mutant proteins (R34W, D337N, and P454L) show aggregates or abnormal leakage-out into cytoplasm (arrowheads) and some WT-like distribution (arrows). Nuclei were stained with DAPI (cyan). (**B-D**) Mutations lowered the neurite arborization activity of CUX2 in fly neurons. Representative images of CUX2-R34W (B) and CUX2-P454L (C). Terminal point numbers were significant decreased in CUX2-P454L, CUX2-W958R and E1283K (n=13 ~ 25) (D). Results of one-way ANOVA followed by Tukey’s test. (**E**) TUNEL assay revealed no apoptosis in adult transgenic flies of CUX2-WT and mutants. Scale bars = 20 μm (A) or 50 μm (B, C, and E). * *P* < 0.05, ** *P* < 0.01, *** *P* < 0.001.

**Figure S2. *Cux2*-deficient mice show seizure susceptibility to kainate but not to PTZ.** (**A**) Body weight was similar among genotypes of 2-month-old mice. (**B-F**) Unchanged seizure susceptibility of *Cux2*-KO mice to PTZ. There are no significant differences in latency to generalized convulsive seizure (GS) (B), latency to death (C), percentage of animals exhibiting GS (D), mortality rate (E), and seizure severity score (F) among genotypes. (**G, H**) *Cux2*-KO mice show increased seizure susceptibility to kainate. Latency to onset of GS (G) and that to death (H) were significantly decreased in *Cux2*(−/−) female and combined gender mice. One-way ANOVA (A-C, F-H) or Pearson’s Chi-square test (3×2 contingency table) (D, E). mean (horizontal bars) ± s.e.m. (B, C, G, H). n or numbers in round brackets =mouse numbers. * *P* < 0.05, ** *P* < 0.01.

**Figure S3. CUX2 antibody specifically recognizes CUX2 protein.** (**A**) Sagittal brain sections from 10-month-old adult WT mice were stained with an antibody to CUX2. Immunosignals (arrows) were observed widely in the brain including in neurons at the hippocampus and cerebral cortex. A2-A5: magnified images outlined in A1. (**B**) CUX2 immunosignals were not observed in *Cux2*(−/−) mouse. B2-B5: magnified images outlined in B1. (**C**) In 2-month-old wild-type mouse, CUX2 (brown) is densely expressed in excitatory neurons at neocortical (II–IV) and entorhinal cortex (II–III) upper layers, but in hippocampal observed in inhibitory but not excitatory neurons. (**D**) In hippocampus at P15, CUX2 (brown) is expressed in interneurons but not in excitatory neurons. Scalebars=500 μm (A1 and B1), 100 μm (C, D) or 20 μm (A2-A5 and B2-B5). CTX; cerebral cortex, DG; dentate gyrus, so; stratum oriens, h; hilus.

**Figure S4. CUX2 is expressed in hippocampal SST-positive, RLN-positive or PV-positive inhibitory neurons.** Tissue sections from P15 (**A-C**) or 2-month-old (**E-G**) WT mouse brains were stained with antibodies to CUX2 (brown) and somatostatin (SST, blue) (A, E), reelin (RLN, blue) (B, F) or parvalbumin (PV, blue) (C, G). CUX2 expression was observed in SST-positive, PV-positive and RLN-positive (arrow) interneurons at both stages. Some of the intense CUX2-positive cells were SST-negative, RLN-negative or PV-negative (white arrow head), and some of the intense SST-positive or PV-positive cells were CUX2-negative (black arrow head). CUX2-positive / PV-positive cell number increased at 2-months compared to P15. A2-A4, B2-B4, C2-C4, E2-E4, F2-F4, G2-G4, A5 and B5: magnified images outlined in A1, B1, C1, E1, F1, G1, A4 and B4, respectively. (**D, H**) Tissue sections from at P15 (D) or 2-month-old (H) WT mouse brain were stained with antibodies to SST (green) or RLN (magenta) and DAPI (cyan). The SST expression was observed in RLN-positive interneurons (arrows) at the hilus of hippocampus. SST/RLN-double positive cells were more frequent in P15 (D) than 2-month-old sections (H). RLN signals became weaker at 2-months (H) compared to P15 (D). Some of intense SST-positive cells were RLN-negative (arrow head), and some of intense RLN-positive cells were SST-negative (asterisk). Scale bar=100 μm (A1, B1, C1, E1, F1, and G1) or 20 μm (A2-A4, B2-B4, C2-C4, D, E2-E4, F2-F4, G2-G4 and H). h; hilus.

**Figure S5. Unchanged cell numbers of hippocampal inhibitory neurons, mossy fibers and cFos expression in *Cux2*-deficient mice.** (**A**) Hippocampus was separated in 7 regions and immunoreactive cells were counted. Cell densities were determined by average number of cells in 4 sections / area. (**B-D**) There were no differences in the number of SST-positive (B), RLN-positive (C) or PV-positive (D) neurons between genotypes. Statistical analyses were performed using one-way ANOVA (*p* > 0.05) (n = WT: 6, *Cux2*(+/−): 6, *Cux2*(−/−): 6). so; stratum oriens, sp; stratum pyramidale, sr; stratum radiatum, lm; lacunosum-moleculare, sm; stratum moleculare, sg; stratum granulosum, h; hilus, w; whole hippocampus. (**E, F**) Timm staining (E) and c-Fos immunohistochemistry (F) did not show differences in *Cux2*-KO mice. Scale bar = 100 μm (A and E).

**Figure S6. IPSCs in PV-positive cells at dentate gyrus were increased, but no differences in EPSCs in CA3 pyramidal cells of *Cux2*-deficient mice.** (**A-D**) In PV-positive cells at dentate gyrus, IPSCs were measured at 6~7-week-old. Amplitude histograms of sIPSC (A) and mIPSC (C) from WT mice and *Cux2*(−/−) mice. Cumulative probability plots and average values (inset) for sIPSC (B) and mIPSC (D) show significant differences in inter-event intervals of sIPSC and mIPSC and amplitude of mIPSC populations derived from *Cux2*(−/−) and significantly higher average of frequency of sIPSC, but unchanged averages of amplitude of sIPSC and both frequency and amplitude of mIPSC in *Cux2*(−/−). (**E-H**) In pyramidal neurons at CA3 region, EPSCs were measured at 6~7-week-old. Amplitude histograms of sEPSC (E) and mEPSC (G) from WT, *Cux2*(+/−) and *Cux2*(−/−) mice. Cumulative probability plots and average values (inset) for sEPSC (F) and mEPSC (H) in WT, *Cux2*(+/−) and *Cux2*(−/−) show no differences in EPSCs among genotypes. (I) No significant difference on MF-evoked EPSCs in pyramidal neurons at CA3 region was observed across genotypes at 6~7-week-old. Statistical analyses were performed using one-way ANOVA followed by Tukey–Kramer Multiple Comparison Test (B, D, F, H, I) or Kolmogorov-Smirnov (K-S) Test (B, D). * *P* < 0.05, *** *P* < 0.001.

**Figure S7. Genome structures of human and mouse *CUX1* and *CASP* in WT and *Casp*-specific knock-out mice.** Exons 1~14 are commonly found in human *CUX1* and *CASP* in human (**A**) and in the corresponding genes in mice (**B**). In *Casp*-KO mice, a mutation (c.1514^1515insTT, p.S506fs) was inserted in Casp-specific exon 17 (**C**). The diagrams were reconstructed from that of UCSC Genome Browser. Vertical lines indicate exons.

**Figure S8. CUX1 is expressed in hippocampal SST-positive, RLN-positive, or PV-positive interneurons.** CUX1 (brown) is expressed in SST-positive (**A**), RLN-positive (**B**) and PV-positive (**C**) interneurons (arrows). Some of intense CUX1-positive cells were SST-negative, RLN-negative or PV-negative (white arrowheads), and some of SST-positive, RLN-positive or PV-positive cells were CUX1-negative (black arrowhead and not shown). Most of CUX1-positive cells were PV-negative (white arrowhead) in hilus (C). A2-A4, B2-B4, C2-C4: magnified images outlined in A1, B1, C1, respectively. Scale bars = 100 μm (A1, B1 and C1) or 20 μm (A2-A4, B2-B4 and C2-C4). sp; stratum pyramidale, so; stratum oriens, sg; stratum granulosum, h; hilus.

**Figure S9. CASP, CUX1 and CUX2 expression and increased seizure susceptibility to kainate with a tendency of increase in excitatory cell number in entorhinal cortex of *Casp*-deficient mice.** (**A**) CASP immunosignals were observed in both excitatory and inhibitory neurons in the hippocampus, though the signals in inhibitory neurons were especially intense such as those at hilus (A3) and stratum oriens of CA3 (A5), while signals in excitatory neurons are tiny such as those at stratum granulosum (A4) or pyramidal of CA3 (A6). CASP signals well disappeared in *Casp*(−/−) mice (A7, A8). Nuclei were stained with hematoxylin (blue). (**B, C**) Unaltered expression and subcellular localization of CUX1 and CUX2 in cerebral cortex (B) and hilus of hippocampus (C) of *Casp*-KO mice. (**D**) Body weights were similar among *Casp*-KO mice and WT littermates at 2-months. (**E**) qPCR experiments showed that mRNA expression levels of kainate receptor subunits were not altered in *Casp*-KO mice at 2-month-old. (**F**) In a Nissl staining, number of entorhinal cortex layer II-III neurons was comparable between WT and *Casp*(−/−) mice (2-month-old). (**G**) Seizure severity scores were significantly higher in *Casp*(−/−) male and combined gender mice. **(H)** Latencies to onset of generalized seizures (GS) were significantly decreased in male and combined gender of *Casp*(−/−) mice. (**I**) Latencies until death were also significantly decreased in combined gender of *Casp*(−/−) mice. Mice without GS or death within 3,600 sec were plotted at "no GS" or "no death", respectively. Circles represent individual mice. One-way ANOVA followed by Tukey–Kramer Multiple Comparison Test (D, E, G-I), or one-way ANOVA test (F). mean (horizontal bars) ± s.e.m. (H, I). n=mouse number. Scale bars= 50 μm (A1, A2, A7, A8, B and C) or 10 μm (A3-A6). * *P* < 0.05, ** *P* < 0.01.

**Figure S10. CASP interacts with CUX2.** mRFP-tagged CUX2 was co-immunoprecipitated with FLAG-tagged CASP. Mutations did not affect the binding. Endophilin: negative control. Original blots are presented in Supplementary Figure S11.

**Figure S11. Full size western blot images.** Original scanned western blot data that were used to generate Figure S10. Dashed rectangles in the images indicate the location of the cropped images. Images of blots with adequate length and membrane edges could not be provided because the blots were cut prior to hybridisation with antibodies and scanned at inside of blots, respectively.

## Notes

### Competing Interest Statement

The authors have declared no competing interest.

## References

1. Zimmer, C., Tiveron, M. C., Bodmer, R. & Cremer, H. Dynamics of Cux2 expression suggests that an early pool of SVZ precursors is fated to become upper cortical layer neurons. Cereb. Cortex 14, 1408–1420 (2004).

2. Cubelos, B. et al. Cux-1 and Cux-2 control the development of Reelin expressing cortical interneurons. Dev. Neurobiol. 68, 917–925 (2008).

3. Cubelos, B. et al. Cux-2 controls the proliferation of neuronal intermediate precursors of the cortical subventricular zone. Cereb. Cortex 18, 1758–1770 (2008).

4. Cubelos, B. et al. Cux1 and Cux2 regulate dendritic branching, spine morphology, and synapses of the upper layer neurons of the cortex. Neuron 66, 523–535 (2010).

5. Suzuki, T. et al. Genome-wide association study of epilepsy in Japanese population identified an associated region at chromosome 12q24. Epilepsia 62, 1391–1400 (2021).

6. Chatron, N. et al. The epilepsy phenotypic spectrum associated with a recurrent CUX2 variant. Ann. Neurol. 83, 926–934 (2018).

7. Barington, M., Risom, L., Ek, J., Uldall, P. & Ostergaard, E. A recurrent de novo CUX2 missense variant associated with intellectual disability, seizures, and autism spectrum disorder. Eur. J. Hum. Genet. 26, 1388–1391 (2018).

8. Téllez-Zenteno, J. F. & Hernández-Ronquillo, L. A review of the epidemiology of temporal lobe epilepsy. Epilepsy Res. Treat. 630853 (2012).

9. Lévesque, M. & Avoli, M. The kainic acid model of temporal lobe epilepsy. Neurosci. Biobehav. Rev. 37, 2887–2899 (2013).

10. Iulianella, A., Sharma, M., Durnin, M., Vanden Heuvel, G. B. & Trainor, P. A. Cux2 (Cutl2) integrates neural progenitor development with cell-cycle progression during spinal cord neurogenesis. Development 135, 729–741 (2008).

11. Gingras, H., Cases, O., Krasilnikova, M., Bérubé, G. & Nepveu, A. Biochemical characterization of the mammalian Cux2 protein. Gene 344, 273–285 (2005).

12. Jinushi-Nakao, S. et al. Knot/Collier and cut control different aspects of dendrite cytoskeleton and synergize to define final arbor shape. Neuron 56, 963–978 (2007).

13. Maher, J. & McLachlan, R. S. Febrile convulsions. Is seizure duration the most important predictor of temporal lobe epilepsy? Brain 118, 1521–1528 (1995).

14. Lievens, P. M., Tufarelli, C., Donady, J. J., Stagg, A., Neufeld, E. J. CASP, a novel, highly conserved alternative-splicing product of the CDP/cut/cux gene, lacks cut-repeat and homeo DNA-binding domains, and interacts with full-length CDP in vitro. Gene 197, 73–81 (1997).

15. Gillingham, A. K., Pfeifer, A. C. & Munro, S. CASP, the alternatively spliced product of the gene encoding the CCAAT-displacement protein transcription factor, is a Golgi membrane protein related to giantin. Mol. Biol. Cell 13, 3761–3774 (2002).

16. Luong, M. X. et al. Genetic ablation of the CDP/Cux protein C terminus results in hair cycle defects and reduced male fertility. Mol. Cell Biol. 22, 1424–1437 (2002).

17. Takata, A. et al. Comprehensive analysis of coding variants highlights genetic complexity in developmental and epileptic encephalopathy. Nat. Commun.10, 2506 (2019).

18. Malsam, J., Satoh, A., Pelletier, L. & Warren, G. Golgin tethers define subpopulations of COPI vesicles. Science 307,1095–1098 (2005).

19. Singh, B. et al. A Kv4.2 truncation mutation in a patient with temporal lobe epilepsy. Neurobiol. Dis. 24, 245–253 (2006).

20. Barnwell, L. F. et al. Kv4.2 knockout mice demonstrate increased susceptibility to convulsant stimulation. Epilepsia 50, 1741–1751 (2009).

21. Li, J. M. et al. Aberrant glutamate receptor 5 expression in temporal lobe epilepsy lesions. Brain Res. 1311, 166–174 (2010).

22. Marksteiner, J., Ortler, M., Bellmann, R. & Sperk, G. Neuropeptide Y biosynthesis is markedly induced in mossy fibers during temporal lobe epilepsy of the rat. Neurosci Lett 112, 143–148.

23. Tonder, N., Kragh, J., Finsen, B. R., Bolwig, T. G. & Zimmer, J. Kindling induces transient changes in neuronal expression of somatostatin, neuropeptide Y, and calbindin in adult rat hippocampus and fascia dentata. Epilepsia 35, 1299–1308.

24. Vezzani, A., Sperk, G. & Colmers, W. F. Neuropeptide Y: emerging evidence for a functional role in seizure modulation. Trends Neurosci 22, 25–30.

25. Bureau, I., Bischoff, S., Heinemann, S. F. & Mulle, C. Kainate receptor-mediated responses in the CA1 field of wild-type and GluR6-deficient mice. J. Neurosci. 19, 653–663 (1999).

26. Fritsch, B., Reis, J., Gasior, M., Kaminski, R. M. & Rogawski, M. A. Role of GluK1 kainate receptors in seizures, epileptic discharges, and epileptogenesis. J. Neurosci. 34, 5765–5775 (2014).

