## Supplementary Information for "CUX2 deficiency causes facilitation of excitatory synaptic transmission onto hippocampus and increased seizure susceptibility to kainate"

Toshimitsu Suzuki[1,2]†, Tetsuya Tatsukawa[2]†, Genki Sudo[2]†, Caroline Delandre[3,18], Yun Jin Pai[3], Hiroyuki Miyamoto[2], Matthieu Raveau[2], Atsushi Shimohata[2,19], Iori Ohmori[4], Shin-ichiro Hamano[5], Kazuhiro Haginoya[6], Mitsugu Uematsu[7], Yukitoshi Takahashi[8,9], Masafumi Morimoto[10], Shinji Fujimoto[11,12], Hitoshi Osaka[13], Hirokazu Oguni[14], Makiko Osawa[14], Atsushi Ishii[15], Shinichi Hirose[15], Sunao Kaneko[16,17], Yushi Inoue[9], Adrian Walton Moore[3], & Kazuhiro Yamakawa[1,2]*

[1]Department of Neurodevelopmental Disorder Genetics, Institute of Brain Science, Nagoya City University Graduate School of Medical Science, Aichi, Japan.

[2]Laboratory for Neurogenetics, RIKEN Center for Brain Science, Saitama, Japan.

[3]Laboratory for Genetic Control of Neuronal Architecture, RIKEN Center for Brain Science, Saitama, Japan.

[4]Department of Special Needs Education, Okayama University Graduate School of Education, Okayama, Japan.

[5]Division of Neurology, Saitama Children's Medical Center, Saitama, Japan.

[6]Department of Pediatric Neurology, Miyagi Children's Hospital, Sendai, Japan.

[7]Department of Pediatrics, Tohoku University School of Medicine, Sendai, Japan.

[8]Department of Pediatrics, Gifu Prefectural Gifu Hospital, Gifu, Japan.

[9]National Epilepsy Center, NHO Shizuoka Institute of Epilepsy and Neurological Disorder, Shizuoka, Japan.

[10]Department of Pediatrics, Kyoto Prefectural University of Medicine, Kyoto, Japan.

[11]Department of Pediatrics, Neonatology and Congenital Disorders, Nagoya City University Graduate School of Medical Sciences, Nagoya, Japan.

[12]Tsutsujigaoka Children's Clinic, Aichi, Japan.

[13]Department of Pediatrics, Jichi Medical University, Shimotsuke, Japan.

[14]Department of Pediatrics, Tokyo Women's Medical University, Tokyo, Japan.

[15]Department of Pediatrics, School of Medicine and Research Institute for the Molecular Pathomechanisms of Epilepsy, Fukuoka University, Fukuoka, Japan.

[16]Department of Neuropsychiatry, Hirosaki University School of Medicine, Hirosaki, Japan.

[17]North Tohoku Epilepsy Center, Minato Hospital, Hachinohe, Japan.

[18]Present address: Menzies Institute for Medical Research, University of Tasmania, Tasmania, Australia.

[19]Present address: Department of Physiology, Nippon Medical School, Tokyo, Japan.

†These authors contributed equally to this work.

* Address correspondence to: Kazuhiro Yamakawa, Ph.D.,

Department of Neurodevelopmental Disorder Genetics, Institute of Brain Science, Nagoya City University Graduate School of Medical Science,

1 Kawasumi, Mizuho-cho, Mizuho-ku Nagoya, Aichi 467-8601, Japan

ORCID iD: https://orcid.org/0000-0002-1478-4390

### Supplemental Methods

#### Quantification of mRNA

For quantification of mRNA expressions in mice and flies, we extracted total RNA from the hippocampus of 2-month-old mice or adult flies using the TRIZOL reagent (Life Technologies) (for flies, six flies for each genotype were extracted together in one tube) and obtained first-strand cDNA from total RNA (1μg) using the Prime Script RT-PCR kit with gDNA Eraser (TaKaRa). Real-time quantitative PCR (RT-qPCR) was performed in triplicate for mice (2-month-old) or in five replicates for adult *Drosophila* using SsoAdvanced Universal SYBR Green Supermix (Bio-Rad) by ABI PRISM 7900. Primers for mice were designed on *Casp* exon17 (forward primer) and exon 18 (reverse primer), *Cux1* exon 23 and 24, and *Cux2* exon 22 and 23. The data were normalized to *Gusb* levels for mice or *Gapdh* levels for *Drosophila*. Data were analyzed by the Delta Delta CT (ΔΔCT) method. Primers are listed in Table S3.

#### Expression constructs and mutagenesis

We amplified the complete open reading frame (ORF) of *CUX2* from the cDNA clone (IMAGE: 8860519, obtained from DNAFORM) and *CASP* from human fetal brain cDNA (Clontech) by PCR using PrimeSTAR HS DNA Polymerase and cloned them into pcDNA3-mRFP (kind gift from Dr. Atsushi Miyawaki, RIKEN CBS, Saitama, Japan) or pcDNA3-FLAG (Invitrogen) (mRFP or FLAG sequence was fused at the N-terminus of the protein of interest). We introduced mutations using the QuickChange Site-Directed Mutagenesis kit (Agilent Technologies) and confirmed nucleotide changes and integrity of the full ORF sequences by DNA sequencing.

#### Cell imaging

We cultured HeLa.S3 cells in 24-well plates containing glass coverslips (12 mm) coated with Poly-L-Lysine (BD Biosciences) and grew them in D-MEM supplemented with 10% FBS, 100 U/mL penicillin, and 100 μg/mL streptomycin. We transfected HeLa.S3 cells with constructs encoding mRFP-tagged wild-type or mutants of CUX2 proteins using Lipofectamine-LTX (Thermo Fisher Scientific). Cultured cells were fixed with 4% paraformaldehyde (PFA, TAAB) in 1X phosphate buffer saline (PBS) 48 h after transfection. Nuclei were stained with 4', 6-diamidino-2-phenylindole (DAPI), dihydrochloride. Images were acquired under a TCS SP2 (Leica) or Biozero BZ-X710 (KEYENCE) microscope. Cells with abnormally localized protein (leakage into the cytoplasm or abnormal aggregates) were determined by manually in a blinded manner. Abnormal aggregates were defined by puncta sizes bigger than the average of WT puncta + 2SD.

**Drosophila stocks and crosses**

*CUX2* constructs for *Drosophila* were made by cloning human *CUX2* (WT or mutants) into pWALIUM10 (Transgenic RNAi Project). The final constructs were inserted into the third chromosome (attP2) using site-specific bacteriophage PhiC31 systems (BestGene Inc.). For ectopic expression experiments, we crossed *UAS-CUX2* to *Gal4*$^{2\text{-}21}$, *UAS-mCD8::GFP* (Jinushi-Nakao *et al*., 2007) and imaged third instar wandering larvae with a Nikon C1 Confocal microscope. Dendritic arbor quantification was carried out using Imaris (Bitplane).

**TUNEL assay in flies**

Adult flies were fixed with 4% PFA in 0.2 M phosphate-buffer (PB) for 1 h. After dehydration by ethanol series (50%, 70%, 90%, 95%, and 100% ethanol), fixed fly samples were treated with 1-Butanol for 1 h. Paraffin (6-μm-thick) sections were used in the assay. Apoptotic cells in *Drosophila* paraffin sections (*Gal4*$^{2\text{-}21}$, *UAS-mCD8::GFP/UAS-CUX2*) and control flies (*Gal4*$^{2\text{-}}$

[21], *UAS-mCD8::GFP*) were detected using the DeadEnd Colorimetric TUNEL System (Promega). For a negative control, sections were incubated in buffer without the recombinant terminal deoxynucleotidyl transferase (rTdT) enzyme. DNase I-treated sections were used as positive controls. Images were acquired by BIOZERO BZ-8100 (KEYENCE).

**Seizure susceptibility in mice**

Seizure susceptibilities of *Cux2*- or *Casp*-KO mice (2 to 3-month-old) to pentylenetetrazole (PTZ, Sigma) or kainate (Sigma) were measured. PTZ or kainate was dissolved in 1X PBS in a total volume of 200–300 μL and administered by intraperitoneal injection (i.p.) at a dose of 50 or 30 mg/kg body weight, respectively. After injections, animals were watched closely for 600 s for PTZ and 3,600 s for kainate. Latencies until onset of myoclonic jerks, tonic seizures, generalized seizure (GS), and death were measured. For mice without myoclonic jerks, tonic seizures, GS, or death, calculation of latency periods of 600 s for PTZ or 3,600 s for kainate were temporarily assigned. For PTZ, seizure severities were graded as follows: stage score 1–myoclonic jerks (twitch), stage score 2–repeated tonic seizures or abortive generalized seizure, stage score 3–fully generalized seizure, stage score 4–tonic-hind limb extension seizure, and stage score 5–death. For kainate, seizure severities were graded as follows: stage score 1–behavioral arrest with mouth/facial movements, stage score 2–head nodding, stage score 3–forelimb clonus, stage score 4–rearing, stage score 5–rearing and falling, stage score 6–loss of posture and generalized convulsive activity, and stage score 7–death. Maximum seizure score for each animal was determined at 600 or 3,600 s after PTZ or kainate administration, respectively. In all experiments, the experimenter was blinded to the animal genotypes.

**Histological analyses**

Mice were perfused intracardially with periodate-lysine-4% PFA solution (PLP; 10 mM $NaIO_4$, 75 mM lysine, 37.5 mM phosphate buffer, with 4% PFA). Paraffin (6-μm-thick) sections were used in most experiments, while frozen (30-μm-thick) sections were used in c-Fos, Zinc transporter-3 (ZnT-3), and Timm staining. Heat-induced epitope retrieval was performed for paraffin sections using a microwave oven in citrate EDTA buffer (10 mM Citric acid, 1 mM EDTA, pH 6.0) for 10 min. The rabbit polyclonal antibody to CUX2 (1:500-1,000 dilution), rabbit polyclonal antibodies to CUX1, M-222 (CDP, 1:500-1,000 dilution, sc13024, Santa Cruz Biotechnology, raised against 1,111-1,332 aa of mouse CUX1) or C-20 (CDP, 1:100 dilution, sc6327, Santa Cruz Biotechnology, raised against C-terminus of mouse CUX1), rabbit monoclonal antibody to CASP (1:100 dilution, EPR18806, ab182216, abcam, antibody raised against 400-650 aa of human CASP but also recognizes mouse and rat), antibody to Somatostatin (SST; 1:1,000 dilution, T-4103, Peninsula Laboratories, LLC), antibody to Reelin (RLN; 1:1,000 dilution, ab78540, G10, abcam), antibody to parvalbumin (PV; 1:500-2,000 dilution, MAB1572, Millipore), antibody to c-Fos (1:5,000 dilution, PC38, Calbiochem), antibody to Doublecortin (1:500 dilution, sc-8066, Santa Cruz Biotechnology), antibody to phospho-Histone H3 (Thr3) (1:2,000 dilution, JY325, 04-746, Millipore), antibody to Ki67 (1:100 dilution, B56, 550609, BD Pharmingen), antibody to NeuN (1:2,000 dilution, XAB377, Millipore), antibody to GFAP (1:2,000 dilution, G3893-2ML, 5-A-G, SIGMA), and antibody to ZnT-3 (1:1,000 dilution, 197002, Synaptic Systems) were used. Secondary antibodies were horse antibody to mouse IgG conjugated to biotin (1:200 dilution, BA-2000, VECTOR Laboratories) or goat antibody to rabbit IgG conjugated to biotin (1:200 dilution, BA-1000, VECTOR Laboratories). For staining using mouse primary antibodies, the Mouse on Mouse (M.O.M.) detection kit (BMK-2202, VECTOR Laboratories) was used to reduce endogenous mouse IgG staining. Immunoreactivity was visualized using a Vectastain Elite ABC kit (VECTOR Laboratories), developed using the ImmPACT DAB Peroxidase (HRP) Substrate kit or VECTOR Blue Alkaline Phosphatase Substrate (VECTOR Laboratories). For fluorescent immunohistochemistry, secondary antibodies were donkey-raised antibodies to mouse IgG conjugated to Alexa Fluor 594 (1:300 dilution, A-21203, Thermo Fisher) or to rabbit IgG conjugated to Alexa Fluor 488 (1:300 dilution, A-21206, Thermo Fisher). Nuclei were stained with DAPI. Normal mouse IgG or rabbit IgG (Santa Cruz Biotechnology) was used as a negative control. Images were acquired by the Biozero BZ-X710 microscope. Timm staining was performed using the FD Rapid TimmStain Kit (FD Neurotechnologies, Inc.). For Nissl staining, 0.25% thionin (pH 3.8) was used. Cell counts in the entorhinal cortex and neocortex were performed manually. Thickness of the CUX1-positive layer was measured using ImageJ (National Institute of Health). Measurements were performed in a blinded manner.

**In vitro electrophysiology**

Hippocampal slices (350 μm thickness) were prepared on a vibratome (PRO 7, DOSAKA EM) under isoflurane anesthesia from female *Cux2*-KO or male *Casp*-specific KO mice (6 to 7-week-old), and incubated at least 1 h at 34°C in equilibrated (95% $O_2$, 5% $CO_2$) artificial CSF (ACSF) containing: 125 mM NaCl, 2.5 mM KCl, 1 mM $MgSO_4$, 2 mM $CaCl_2$, 1.25 mM $NaH_2PO_4$, 26 mM $NaHCO_3$, 11 mM glucose, 3 mM Na-pyruvate, and 1 mM Na-L-ascorbic acid before recording. Whole-cell patch-clamp recordings were performed at 30–31°C using a MultiClamp-700B amplifier and pClamp 10.3 software (Molecular Devices). Patch pipettes (4–6 MΩ) were pulled from borosilicate glass on a puller (DMZ-UNIVERSAL PULLER, Zeitz Instruments). EPSCs or IPSCs were recorded using electrodes filled with an internal solution containing: 128 mM Cs-methansulfate, 6 mM KCl, 2 mM NaCl, 0.2 mM EGTA, 20 mM HEPES, 4 mM

MgATP, 0.3 mM $Na_3GTP$, 14 mM phosphocreatine, and 5 mM QX314. Data were discarded when series resistance varied by more than 20%. To investigate perforant path synaptic transmission, perforant path-evoked EPSCs (eEPSCs) were recorded from dentate granule cells. Constant current pulses (40 µs duration, 10–100 µA amplitude, every 15 s 10 times) were applied on the perforant path at the border between the entorhinal cortex and subiculum. To investigate the effects of kainate to inhibit dentate granule cells, spontaneous IPSCs were recorded from dentate granule cells held at 0 mV in the presence of NMDA receptor antagonist D-AP5 (50 µM, Tocris Bioscience) and AMPA receptor antagonist GYKI 52466 (40 µM, Tocris Bioscience). To investigate the effects of kainate on mossy-fiber (MF) eEPSCs, MF- EPSCs that showed paired pulse facilitation were monitored from CA3 pyramidal cells held at -70 mV by stimulation through a bipolar electrode placed in the striatum lucidum. To monitor miniature EPSCs, 1 µM tetrodotoxin (TTX, Tocris Bioscience) was added to ACSF. Picrotoxin (100 µM, Sigma) and strychnine (1 µM, Sigma) are routinely used in the ACSF to block inhibitory inputs. Data were filtered (1 kHz), digitized (10 kHz), stored, and analyzed using Clampfit 10.3 (Molecular Devices), Origin 8.5J (OriginLab Corporation), and Minianalysis software (Synaptosoft, Decatur).

**Co-immunoprecipitation**

Subconfluent 293T cells were cotransfected with pcDNA3-mRFP-CUX2 and pcDNA3-FLAG-CASP or pcDNA3-FLAG-Endophilin as a control using Lipofectamine 2000. At 20 h after transfection, cells were lysed in 10 mM Tris (pH 8.0), 150 mM NaCl, 5 mM EDTA, 0.5% NP-40, and protease inhibitor (Complete, Roche). FLAG antibody-Agarose (Sigma) was added into the lysate and incubated overnight at 4°C. Next, protein complexes were washed 5 times with 1 mL lysis buffer and resolved by 4%–20% SDS-PAGE, after which Western blots were performed with DsRed antibody (Clontech). For CUX2 detection, Western blots were performed using the FLAG antibody (Sigma).

**URLs**

Simple Modular Architecture Research Tool (SMART), http://smart.embl-heidelberg.de; The Human Genetic Variation Database (J-HGVD), http://www.genome.med.kyoto-u.ac.jp/SnpDB; Exome Variant Server (EVS), NHLBI GO Exome Sequencing Project (ESP), Seattle, WA, http://evs.gs.washington.edu/EVS/; 1000 Genomes Browser (1kGP), http://www.ncbi.nlm.nih.gov/variation/tools/1000genomes; Exome Aggregation Consortium (ExAC), http://exac.broadinstitute.org; Genome Aggregation Database (gnomAD v2.1.1), https://gnomad.broadinstitute.org; Polyphen-2, http://genetics.bwh.harvard.edu/pph2/; Mutation taster, http://mutationtaster.org/; PROVEAN, http://provean.jcvi.org/about.v1.0.php; SIFT, http://sift.jcvi.org/; M-CAP, http://bejerano.stanford.edu/mcap/; UCSC Genome Browse, http://genome.ucsc.edu/index.html; TIGM, http://www.tigm.org/; BestGene Inc., http://www.thebestgene.com; Addgene, http://www.addgene.org; Feng Zhang lab's Target Finder, http://crispr.mit.edu.

**Reference for Supplemental Methods**

Jinushi-Nakao S, Arvind R, Amikura R, Kinameri E, Liu AW, and Moore AW. Knot/Collier and cut control different aspects of dendrite cytoskeleton and synergize to define final arbor shape. Neuron 2007;56:963–78.

### Supplemental Note

#### Disease phenotypes of patients with *CUX2*, *CUX1* or *CASP* mutations

SIZ-004

A 37-year-old female without family history for epilepsy had febrile convulsions several times per year from age 6 months until 4 years. At age 6 years, she had a convulsion shortly after a traffic accident without any neurological sequelae. At age 11, she started to have unaware focal seizures with behavioral automatism, which repeated monthly despite treatment, and rare convulsions. She had no warning sign before loss of awareness. She had a right ICPC aneurysm, for which a clipping surgery was performed. EEG showed interictal spikes and ictal slow waves on the right temporal region. MRI showed right hippocampal atrophy. Her WAIS-IQ was 56. She underwent right selective amygdalohippocampectomy and became seizure-free. The pathology was mesial temporal sclerosis.

SIZ-014

A 42-year-old male had a febrile convulsive status epilepticus lasting 10 hours at age 1 year 2 months with residual right hemiparesis for 1 month. He also suffered from nephrotic syndrome in his early childhood. His father's sister suffered from epilepsy. Beginning at age 3, he sometimes complained about abdominal pain, which, after age 10, evolved into loss of consciousness with right hand dystonia and oral automatism occurring weekly. EEG showed spikes in the temporal area predominantly on the left side and left side starting ictal pattern. MRI revealed left hippocampal atrophy with increased signal. He was left handed and his IQ was 74.

He underwent selective amygdalohippocampectomy on the eft side and has only sporadic abdominal pain post surgically. The pathology was mesial temporal sclerosis.

SIZ-016

A 24-year-old male without relevant family or personal history had a sudden headache and a generalized convulsion at age 15. An MRI revealed a hemorrhagic region in the right temporal lobe as well as right hippocampal atrophy. One month later he started to experience brief epigastric discomfort weekly. Since 20 years of age, this discomfort began to be followed by staring, lip smacking, and hypersalivation as well as left hand dystonia. His EEG showed sharp waves in the right temporal region interictally and rhythmic discharges starting at right frontotemporal electrodes ictally. MEG also showed clustering of dipoles in the right temporal lobe. His WAIS-IQ was 91. As the seizure was intractable to medication, he underwent right temporal lobectomy at 24 years of age. The pathology was mesial temporal sclerosis.

SIZ-022

A 29-year-old female had febrile convulsive status at the age of 1 year with a sequela of hemiparesis. She started to have unaware focal seizures every month since the age of 8 and noticed an aura since the age of 15. EEG and MRI suggested medial temporal lobe epilepsy. Her WAIS-IQ was 89. She underwent selective amygdalohippocampectomy on the left side at age 29 years, and the pathology was mesial temporal sclerosis.

SIZ-060

This 26-year old male had 6 febrile convulsions between ages 1 and 3. At age 17 he had his first spell of loss of awareness with increased tonicity and oral automatism after feeling an unpleasant sensation. The seizure repeated weekly. The EEG showed bilateral independent sharp waves in the temporal regions interictally and rhythmic slow waves starting on the left side ictally. MRI revealed increased intensity in the left mesial temporal lobe. His WAIS-IQ was 64. He underwent standard anterior temporal lobectomy at age 24 years and became seizure free. The pathology was mesial temporal sclerosis.

SIZ-063

A 9-year-old girl had a febrile convulsion at 5 years of age. There was a history of remote febrile convulsion on her mother's side. Absence seizure with bland facial expression started at the age of 8 and increased in frequency; hence, she was administered valproate. The seizures disappeared immediately and completely. EEG showed generalized spike-waves. She was diagnosed with childhood absence epilepsy.

SIZ-068

This 28-year old female had myoclonic and bilateral convulsive seizures since the age of 12 years. Myoclonus was induced by praxis activity but not by photic stimuli. EEG showed generalized spike-waves. There was no family history of epilepsy and no relevant personal history. He is on valproate and is seizure free.

SIZ-073

A 27-year-old male suffered from poliomyelitis at age 1 year 3 months resulting in right leg monoparesis. He had febrile convulsions more than 10 times until 6 years of age. At age 19, he started to have unaware focal seizures consisting of mumbling or oral automatism and stiffening of the left side as well as right hand automatism repeating monthly. He had feelings of anxiety at the onset of seizure. In addition, convulsive seizure with the eyes and head turning towards the left occurred 1–2 times a year. At age 45, he had status epilepticus lasting 2 hours. EEG showed frequent spikes in the right temporal region interictally and ictal discharges starting in the right temporal region. MRI showed right hippocampal atrophy with signal change. His WAIS-IQ was 86. He underwent selective amygdalohippocampectomy on the right side. The pathology was mesial temporal sclerosis.

SIZ-079

A 27-year old male suffered head trauma due to a traffic accident at age 4, but he had no loss of consciousness and not were any abnormalities detected in the CT scan. Since 13 years of age, he started to have seizures yearly, which consisted of tinnitus on the left side and twitching movement of the left half of the face followed by a bilateral convulsive seizure. He experienced numbness of the left upper extremity postictally. EEG showed no epileptiform discharges. There was no apparent lesion on MRI. The seizure disappeared by introducing carbamazepine.

SIZ-127

A 31-year old female had myoclonic seizures since the age of 14 and 2 GTCs at age 15. Valproate administration stopped her seizures. She stopped VPA at age 22, but myoclonus reappeared at age 25 and a GTC at age 26, so she restarted VPA and has had no seizure since then. EEG and MRI were unremarkable. Her brother had a febrile convulsion. She was diagnosed with juvenile myoclonic epilepsy.

SIZ-190

This 34-year-old female had several febrile convulsions before 4 years of age. She started to have epigastric aura and subsequent verbal automatism, staring and oral automatism, which repeated weekly despite medication. EEG showed bilateral independent sharp waves in the temporal regions, but more on the left side. Ictal discharges started more often on the right side. MRI revealed smaller hippocampus on the right side with increased intensity. Her WAIS-IQ was 85. She underwent right amygdalohippocampectomy. The pathology was mesial temporal sclerosis.

SIZ-220

A 27-year-old male had a febrile convulsion at age 1 year. His brother also had a febrile convulsion. The patient started to have unaware focal seizures at age 12 years which disappeared soon after taking medication but relapsed 1 year after stopping medication (age 18 years). The seizure then became intractable. The seizure consisting of staring, oral and verbal automatism occurring weekly was preceded by fearful feeling. Interictal and ictal EEG suggested right temporal seizure focus. MRI revealed right hippocampal atrophy. His WAIS-IQ was 90. He underwent right anterior temporal lobectomy at age 21 years resulting in seizure freedom. The pathology was mesial temporal sclerosis.

SIZ-296

A 15-year-old male had normal development. He started to have astatic seizures and clonic seizures since the age of 2 years. Astatic seizure caused falling with injuries. EEG showed frequent bilateral diffuse irregular spike-waves. MRI was unremarkable. The medications, including valproate, suppressed these seizures completely and the treatment was stopped at age 15. He was diagnosed with epilepsy with myoclonic-atonic seizures (Doose syndrome).

SIZ-456

A 5-year-old girl had febrile and afebrile convulsions since age 1 year. At age 7 years she had brief spells of unconsciousness (absence seizures) occurring every day. EEG showed spike-waves of 3 Hz. She started to take valproate and then has no seizures. There was no family history of epilepsy. She was diagnosed with childhood absence epilepsy.

SIZ-575

A 5-year old girl had no family history and normal development before the onset of epilepsy at age 2 years 1 month when she had afebrile convulsive status epilepticus lasting 40 minutes until receiving diazepam injection. At age 3 years 11 months she had clonic seizure with forward flexion which rapidly increased in frequency till more than 200 per day. Clonazepam abolished this condition but recurred after 3 months. After trials of several AEDs zonisamide finally stopped the seizure. EEG showed frequent bilateral diffuse spike-waves, and MEG showed dipoles localized in the right frontal area. Although MRI was normal, ictal SPECT revealed hyperperfusion on the right frontal area. Developmental quotient was 64 at age 4 years. She was suspected to have frontal lobe epilepsy.

SIZ-638

A 48-year-old male had a GTC at age 15 years and unaware focal seizures without aura occurring weekly since age 33 years. There was no family history of epilepsy. EEG showed abnormal discharges in the right temporal region, though there was no MRI abnormality. After intracranial EEG evaluation, he underwent right amygdalohippocampectomy. The seizure did not disappear completely. There was no pathological change in the resected specimen.

SIZ-669

This 30-year old female has, since shortly after birth, startle-induced myoclonic or tonic seizure followed by bilateral convulsion with sometimes resulting in postictal hemiparesis on the right side. The seizure occurred frequently and was very intractable. EEG showed left frontotemporal dominant spikes and runs of spikes. Sensory-evoked potential study was normal, and her WAIS-IQ was 50. MRI was unremarkable. Her cousin had febrile convulsions.

SIZ-784

A 32-year old male had absences, myoclonus, and rare bilateral convulsive seizures since the age of 16 years. He sometimes fell down and dropped out the things he was holding. His EEG showed bilateral spike-waves without photosensitivity. Sensory-evoked potential study revealed no abnormality. He was seizure-free after valproate administration.

SIZ-891

This 8-year old male had fever-induced convulsive seizures repeated yearly since age 7 months, and afebrile convulsive seizures at age 6 and 7 years. His grand cousin also had febrile convulsion. The EEG and MRI results were unremarkable. He has had no seizure since the administration of valproate was started. In addition, he has autism spectrum disorder.

**Table S1: Number of patients per diagnosis.**

| Sample set | Etiology of epilepsy | Diagnosis | Number of patients |
|---|---|---|---|
| | | CAE | 10 |
| | | GE | 3 |
| | | GEFS+ | 10 |
| | | IGE | 8 |
| | | IGE with myclonis seizure | 1 |
| | | JAE | 10 |
| | Genetic generalized | JME | 61 |
| | | JME closely related | 2 |
| | | Suspected CAE | 2 |
| | | Suspected GEFS+ | 5 |
| | | Suspected IGE | 1 |
| | | Suspected JME | 3 |
| | | Coppora | 1 |
| | | Doose syndrome | 5 |
| | | Doose symptom closely related | 1 |
| | | Epilepsy, ASD | 5 |
| 1st set (271 samples) | | FLE | 4 |
| | | GE | 2 |
| | | Hyperthermia induced partial epilepsy | 2 |
| | | ICEGTC | 4 |
| | | ICEGTC closely related | 1 |
| | | IE | 1 |
| | | Landau-Kleffner syndrome | 1 |
| | Structural/metabolic | lTLE | 11 |
| | | MAE | 2 |
| | | mTLE | 57 |
| | | OLE | 2 |
| | | PE | 29 |
| | | SGE | 15 |
| | | SMEB | 2 |
| | | SMEI | 4 |
| | | Suspected FLE | 1 |
| | | Suspected PE | 1 |
| | | Suspected SMEI | 4 |
| Additional TLE set (69 samples) | | mTLE | 4 |
| | Structural/metabolic | TLE | 62 |
| | | Suspected mTLE | 3 |

CAE: childhood absence epilepsy, GE: generalized epilepsy, GEFS+: generalized epilepsy with febrile seizure plus, IGE: idiopathic generalized epilepsy, JAE: juvenile absence epilepsy, JME: juvenile myoclonic epilepsy, ASD: autism spectrum disorder, FLE: frontal lobe epilepsy, ICEGTC: intractable childhood epilepsy with generalized tonic clonic seizures, IE: intractable epilepsy, lTLE: lateral temporal lobe epilepsy, MAE: myoclonic atonic epilepsy, mTLE: mesial temporal lobe epilepsy, OLE: Occipital Lobe Epilepsy, PE: partial epilepsy, SGE: symptomatic generalized epilepsy, SMEI: severe myoclonic epilepsy in infancy.

**Table S2: Ratio of CUX2-positive cells in hippocampal interneuron subtypes at 2-months of age.**

| | % of CUX2+ cells | Number of CUX2+ cells | Number of cells / section |
|---|---|---|---|
| SST+ | 55.2 | 180 | 29.6 |
| PV+ | 34.1 | 110 | 20.2 |
| RLN+ | 24.9 | 125 | 55.8 |

Number of CUX2+ interneurons were counted in hippocampus of WT mice (n=3) (n=SST+ cell: 326 cells in 11 sections, PV+ cell: 323 cells in 16 sections and RLN+ cell: 502 cells in 9 sections).

**Table S3: Primers used for gene expression analyses of mouse and fly.**

| Gene | Name | Sequence (5' to 3') |
|---|---|---|
| *GluK1* | forward primer | CACGAGACGGCTGCTGAA |
| | reverse primer | ACCACTGTACCTGTAGAGTTCCA |
| *GluK2* | forward primer | ACAATCAACAGGAACAGGACTCT |
| | reverse primer | TGCTGATGAACTGTGTGAAGGA |
| *GluK3* | forward primer | GCTCAGAGGTGGTGGAGAATA |
| | reverse primer | GCCGTGTAGGAGGAGATGAT |
| *GluK4* | forward primer | CGCATGGTAGAATTGGAAGGT |
| | reverse primer | AAGAGACTGTCAGAGATGTTGGA |
| *GluK5* | forward primer | CCACCTTGTCCTCCGTAA |
| | reverse primer | CTCCACGATACCATCCAGAT |
| *Cux2* | forward primer | AGAAGGAGGCTCTGCGGAAG |
| | reverse primer | CATTTCACGGCGCATCCTGG |
| *Cux1* | forward primer | CAGCACCGTCATCAACTGGTTC |
| | reverse primer | AGCTGAAGGTGAGTCGCTGG |
| *Casp* | forward primer | TCCATCCAACGGCCTGATGC |
| | reverse primer | TGATGTTGTCAGCGCGCAGG |
| *Gusb* | forward primer | ACTGACACCTCCATGTATCCCAAG |
| | reverse primer | CAGTAGGTCACCAGCCCGATG |
| *CUX2* | forward primer | TGTGGCTCTCTGACCAGCTC |
| | reverse primer | AGCTCTTCTCAGGCTCTGTGG |
| *Dm Gapdh* | forward primer | TACAGCCCCGACATGAAGGT |
| | reverse primer | GTCATCAGACCCTCGACGATCT |

Dm; Drosophila melanogaster

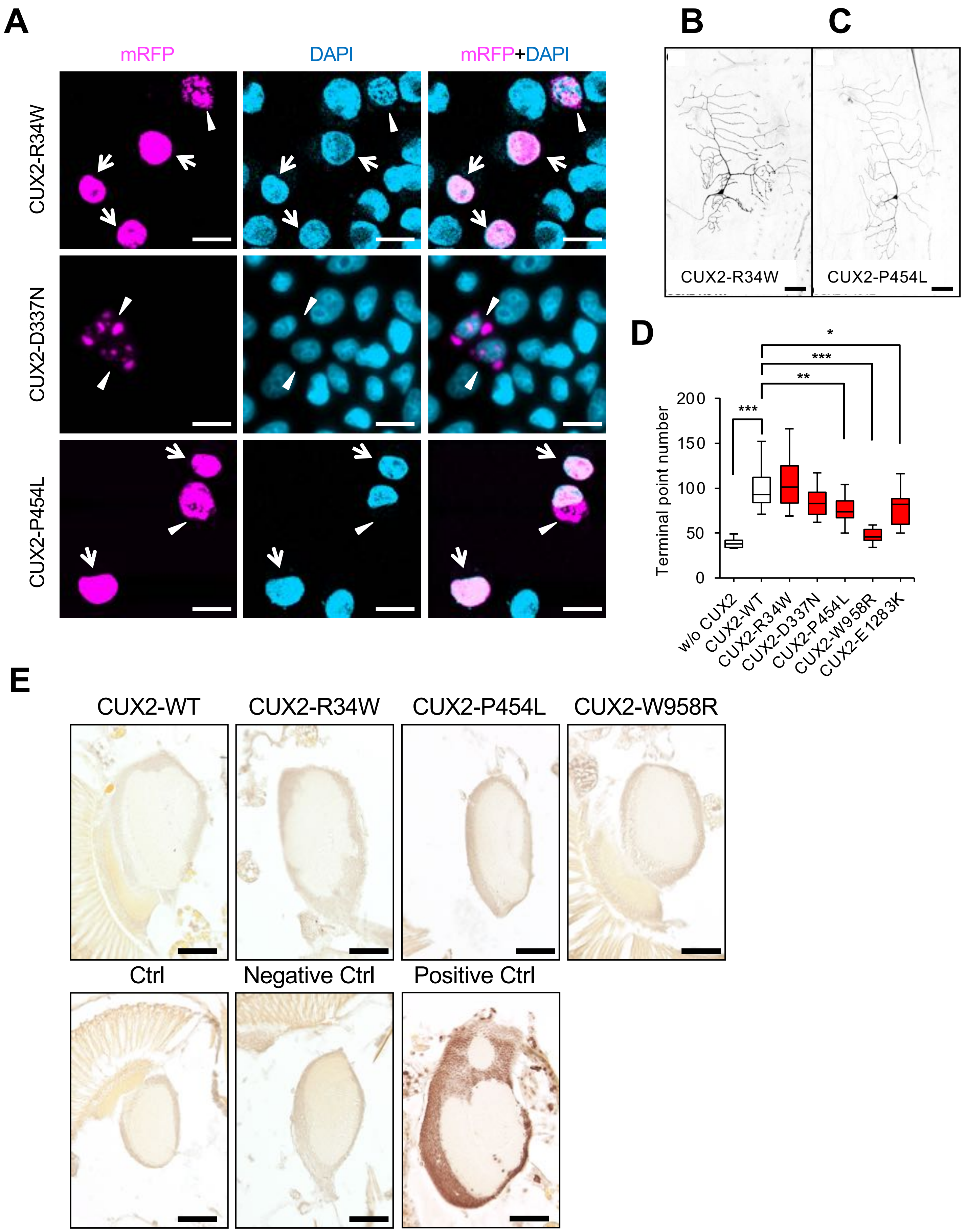


**Figure S1. CUX2 mutants show abnormal subcellular localizations in human cultured cells and decreased branching effects but no apoptosis in fly.** (**A**) In HeLa.S3 cells, CUX2 mutant proteins (R34W, D337N, and P454L) show aggregates or abnormal leakage-out into cytoplasm (arrowheads) and some WT-like distribution (arrows). Nuclei were stained with DAPI (cyan). (**B-D**) Mutations lowered the neurite arborization activity of CUX2 in fly neurons. Representative images of CUX2-R34W (B) and CUX2-P454L (C). Terminal point numbers were significant decreased in CUX2-P454L, CUX2-W958R and E1283K (n=13 ~ 25) (D). Results of one-way ANOVA followed by Tukey's test. (**E**) TUNEL assay revealed no apoptosis in adult transgenic flies of CUX2-WT and mutants. Scale bars = 20 μm (A) or 50 μm (B, C, and E). * $P < 0.05$, ** $P < 0.01$, *** $P < 0.001$.

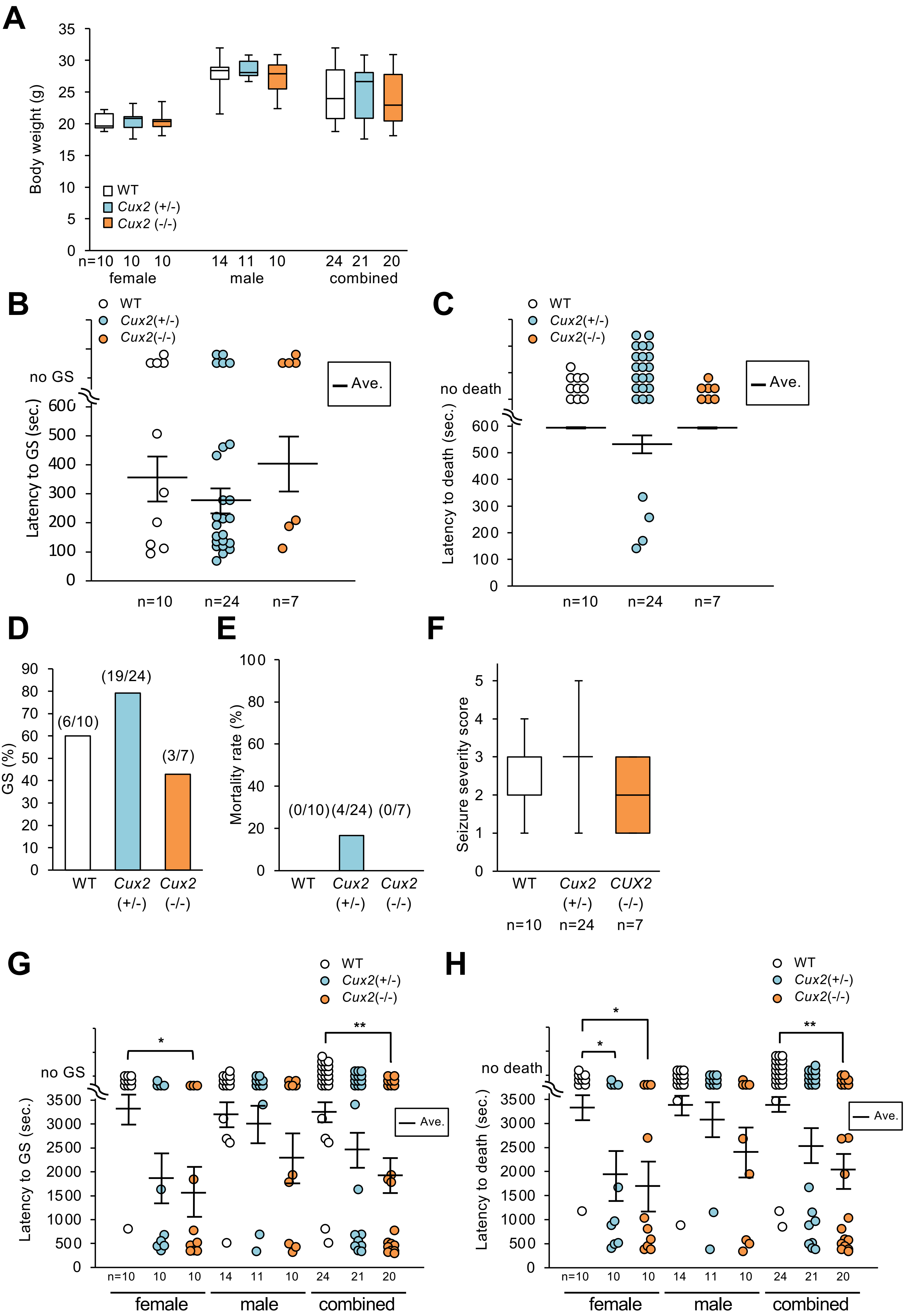
A
Body weight (g)
WT
Cux2 (+/-)
Cux2 (-/-)
n=10 10 10
female
14 11 10
male
24 21 20
combined
B
WT
Cux2(+/-)
Cux2(-/-)
no GS
Latency to GS (sec.)
Ave.
n=10 n=24 n=7
C
WT
Cux2(+/-)
Cux2(-/-)
no death
Latency to death (sec.)
Ave.
n=10 n=24 n=7
D
GS (%)
(6/10)
(19/24)
(3/7)
WT
Cux2 (+/-)
Cux2 (-/-)
E
Mortality rate (%)
(0/10) (4/24) (0/7)
WT
Cux2 (+/-)
Cux2 (-/-)
F
Seizure severity score
WT
Cux2 (+/-)
CUX2 (-/-)
n=10 n=24 n=7
G
WT
Cux2(+/-)
Cux2(-/-)
no GS
Latency to GS (sec.)
Ave.
*
**
n=10 10 10 14 11 10 24 21 20
female
male
combined
H
WT
Cux2(+/-)
Cux2(-/-)
no death
Latency to death (sec.)
Ave.
*
*
**
n=10 10 10 14 11 10 24 21 20
female
male
combined

**Figure S2. *Cux2*-deficient mice show seizure susceptibility to kainate but not to PTZ. (A)** Body weight was similar among genotypes of 2-month-old mice. (**B-F**) Unchanged seizure susceptibility of *Cux2*-deficient mice to PTZ. There are no significant differences in latency to generalized convulsive seizure (GS) (B), latency to death (C), percentage of animals exhibiting GS (D), mortality rate (E), and seizure severity score (F) among genotypes. (**G, H**) *Cux2*-deficient mice show increased seizure susceptibility to kainate. Latency to onset of GS (G) and that to death (H) were significantly decreased in *Cux2*(-/-) female and combined gender mice. One-way ANOVA (A-C, F-H) or Pearson's Chi-square test (3×2 contingency table) (D, E). mean (horizontal bars) ± s.e.m. (B, C, G, H). n or numbers in round brackets =mouse numbers. * $P < 0.05$, ** $P < 0.01$.

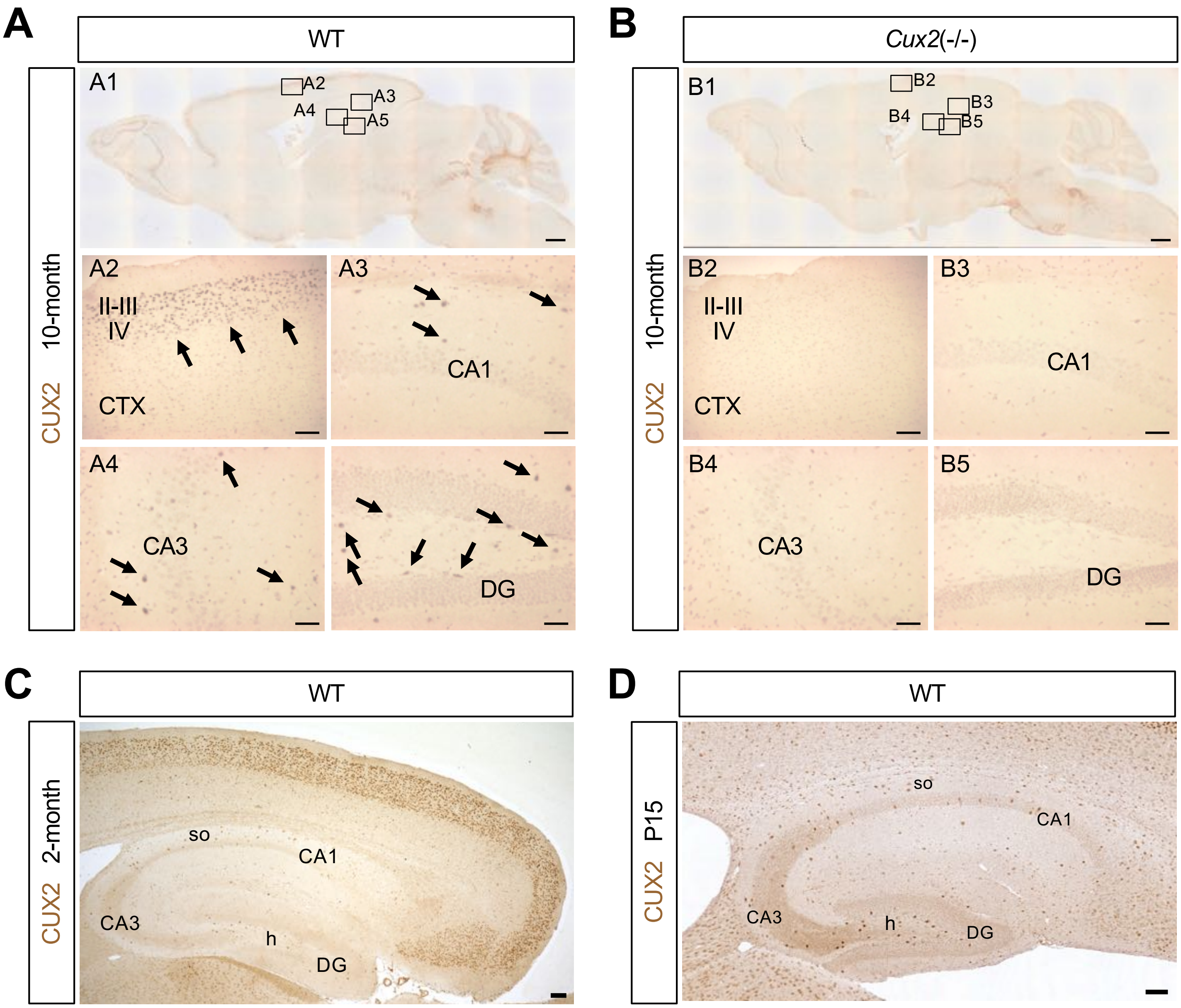


**Figure S3. CUX2 antibody specifically recognizes CUX2 protein.** **(A)** Sagittal brain sections from 10-month-old adult WT mice were stained with an antibody to CUX2. Immunosignals (arrows) were observed widely in the brain including in neurons at the hippocampus and cerebral cortex. A2-A5: magnified images outlined in A1. **(B)** CUX2 immunosignals were not observed in *Cux2*(-/-) mouse. B2-B5: magnified images outlined in B1. **(C)** In 2-month-old wild-type mouse, CUX2 (brown) is densely expressed in excitatory neurons at neocortical (II–IV) and entorhinal cortex (II–III) upper layers, but in hippocampal observed in inhibitory but not excitatory neurons. **(D)** In hippocampus at P15, CUX2 (brown) is expressed in interneurons but not in excitatory neurons. Scale bars=500 µm (A1 and B1), 100 µm (C, D) or 20 µm (A2-A5 and B2-B5). CTX; cerebral cortex, DG; dentate gyrus, so; stratum oriens, h; hilus.

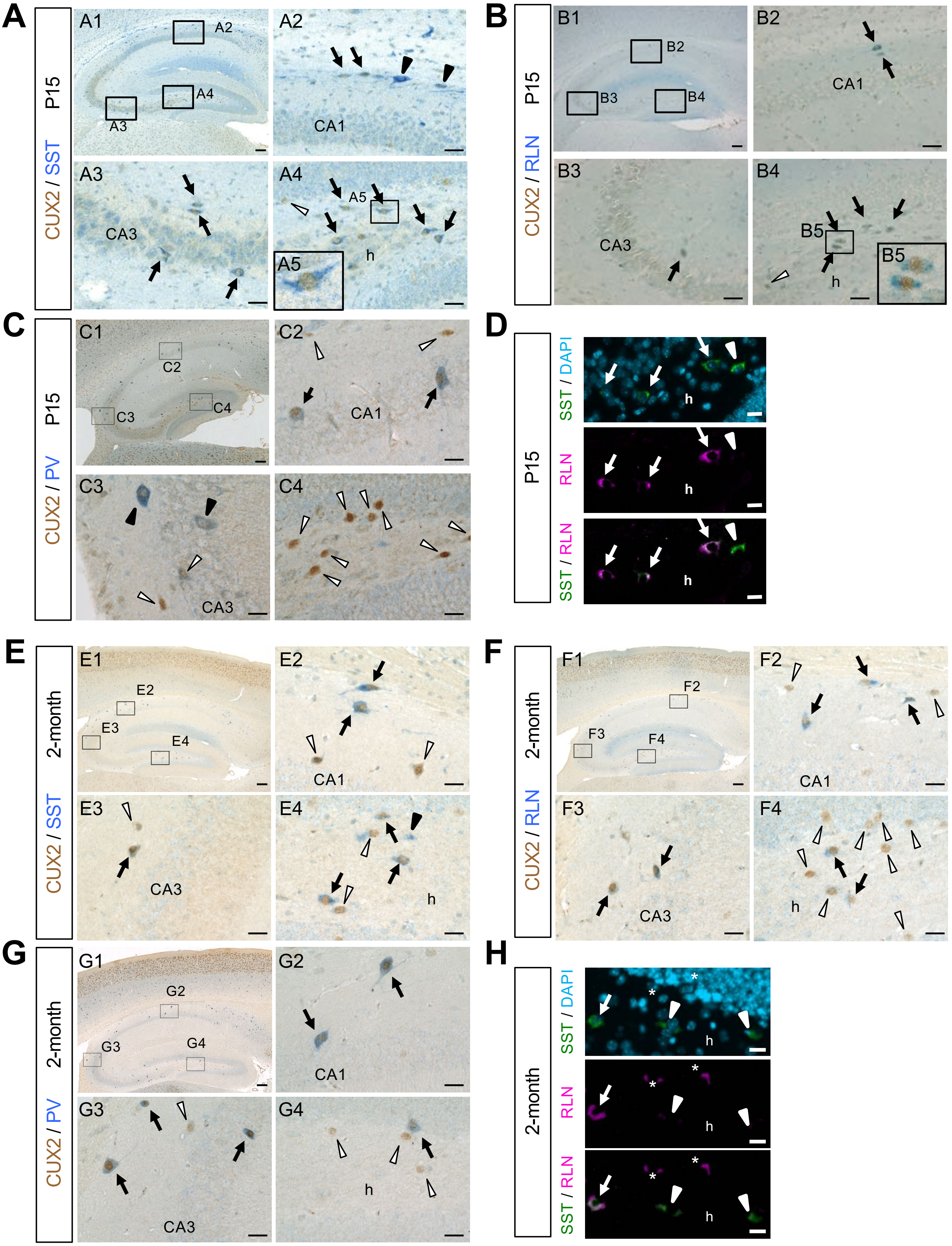

A
P15
CUX2 / SST
A1
A2
A3
A4
A5
CA1
CA3
h
B
P15
CUX2 / RLN
B1
B2
B3
B4
B5
CA1
CA3
h
C
P15
CUX2 / PV
C1
C2
C3
C4
CA1
CA3
D
P15
SST / DAPI
RLN
SST / RLN
h
E
2-month
CUX2 / SST
E1
E2
E3
E4
CA1
CA3
h
F
2-month
CUX2 / RLN
F1
F2
F3
F4
CA1
CA3
h
G
2-month
CUX2 / PV
G1
G2
G3
G4
CA1
CA3
h
H
2-month
SST / DAPI
RLN
SST / RLN
h

**Figure S4. CUX2 is expressed in hippocampal SST-positive, RLN-positive or PV-positive inhibitory neurons.** Tissue sections from P15 **(A-C)** or 2-month-old **(E-G)** WT mouse brains were stained with antibodies to CUX2 (brown) and somatostatin (SST, blue) (A, E), reelin (RLN, blue) (B, F) or parvalbumin (PV, blue) (C, G). CUX2 expression was observed in SST-positive, PV-positive and RLN-positive (arrow) interneurons at both stages. Some of the intense CUX2-positive cells were SST-negative, RLN-negative or PV-negative (white arrow head), and some of the intense SST-positive or PV-positive cells were CUX2-negative (black arrow head). CUX2-positive / PV-positive cell number increased at 2-months compared to P15. A2-A4, B2-B4, C2-C4, E2-E4, F2-F4, G2-G4, A5 and B5: magnified images outlined in A1, B1, C1, E1, F1, G1, A4 and B4, respectively. (**D, H**) Tissue sections from at P15 (D) or 2-month-old (H) WT mouse brain were stained with antibodies to SST (green) or RLN (magenta) and DAPI (cyan). The SST expression was observed in RLN-positive interneurons (arrows) at the hilus of hippocampus. SST/RLN-double positive cells were more frequent in P15 (D) than 2-month-old sections (H). RLN signals became weaker at 2-months (H) compared to P15 (D). Some of intense SST-positive cells were RLN-negative (arrow head), and some of intense RLN-positive cells were SST-negative (asterisk). Scale bar=100 µm (A1, B1, C1, E1, F1, and G1) or 20 µm (A2-A4, B2-B4, C2-C4, D, E2-E4, F2-F4, G2-G4 and H). h; hilus.

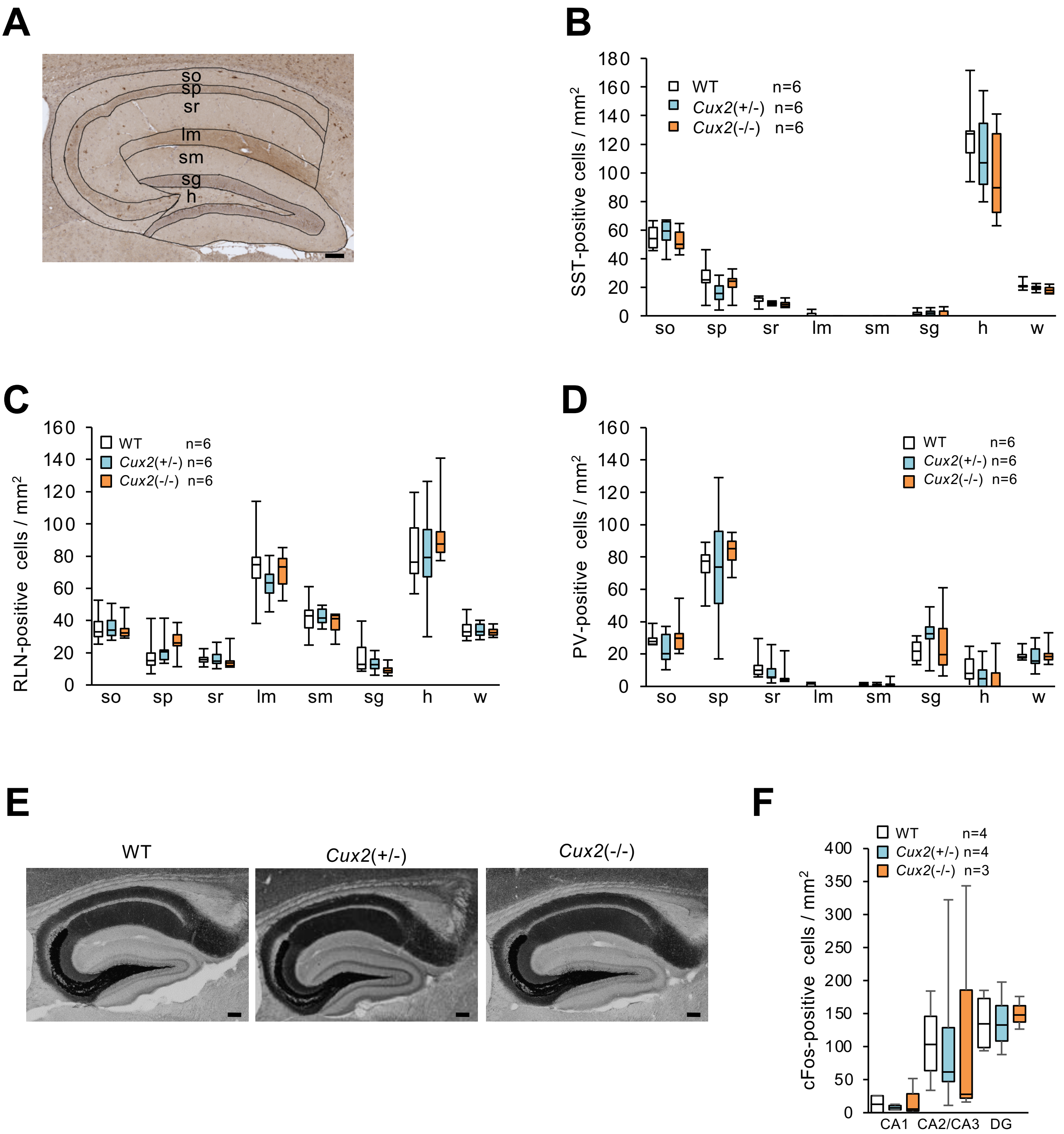


**Figure S5. Unchanged cell numbers of hippocampal inhibitory neurons, mossy fibers and cFos expression in *Cux2*-deficient mice.** (**A**) Hippocampus was separated in 7 regions and immunoreactive cells were counted. Cell densities were determined by average number of cells in 4 sections / area. (**B-D**) There were no differences in the number of SST-positive (B), RLN-positive (C) or PV-positive (D) neurons between genotypes. Statistical analyses were performed using one-way ANOVA ($p > 0.05$) (n = WT: 6, *Cux2*(+/-): 6, *Cux2*(-/-): 6). so; stratum oriens, sp; stratum pyramidale, sr; stratum radiatum, lm; lacunosum-moleculare, sm; stratum moleculare, sg; stratum granulosum, h; hilus, w; whole hippocampus. (**E, F**) Timm staining (E) and c-Fos immunohistochemistry (F) did not show differences in *Cux2*-deficient mice. Scale bar = 100 µm (A and E).

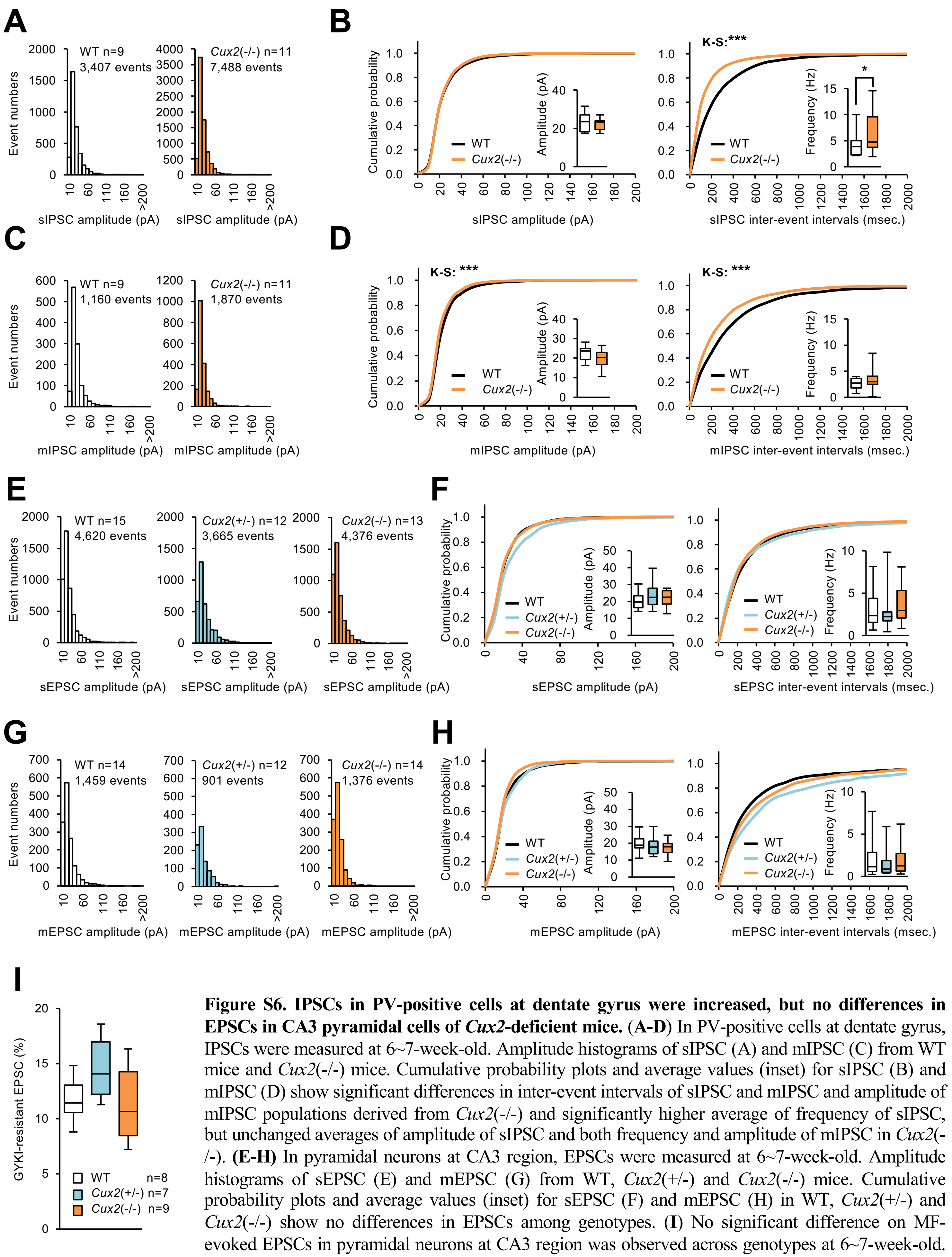

**Figure S6. IPSCs in PV-positive cells at dentate gyrus were increased, but no differences in EPSCs in CA3 pyramidal cells of *Cux2*-deficient mice.** **(A-D)** In PV-positive cells at dentate gyrus, IPSCs were measured at 6~7-week-old. Amplitude histograms of sIPSC (A) and mIPSC (C) from WT mice and *Cux2*(-/-) mice. Cumulative probability plots and average values (inset) for sIPSC (B) and mIPSC (D) show significant differences in inter-event intervals of sIPSC and mIPSC and amplitude of mIPSC populations derived from *Cux2*(-/-) and significantly higher average of frequency of sIPSC, but unchanged averages of amplitude of sIPSC and both frequency and amplitude of mIPSC in *Cux2*(-/-). **(E-H)** In pyramidal neurons at CA3 region, EPSCs were measured at 6~7-week-old. Amplitude histograms of sEPSC (E) and mEPSC (G) from WT, *Cux2*(+/-) and *Cux2*(-/-) mice. Cumulative probability plots and average values (inset) for sEPSC (F) and mEPSC (H) in WT, *Cux2*(+/-) and *Cux2*(-/-) show no differences in EPSCs among genotypes. **(I)** No significant difference on MF-evoked EPSCs in pyramidal neurons at CA3 region was observed across genotypes at 6~7-week-old. Statistical analyses were performed using one-way ANOVA followed by Tukey–Kramer Multiple Comparison Test (B, D, F, H, I) or Kolmogorov-Smirnov (K-S) Test (B, D). * $P < 0.05$, *** $P < 0.001$.

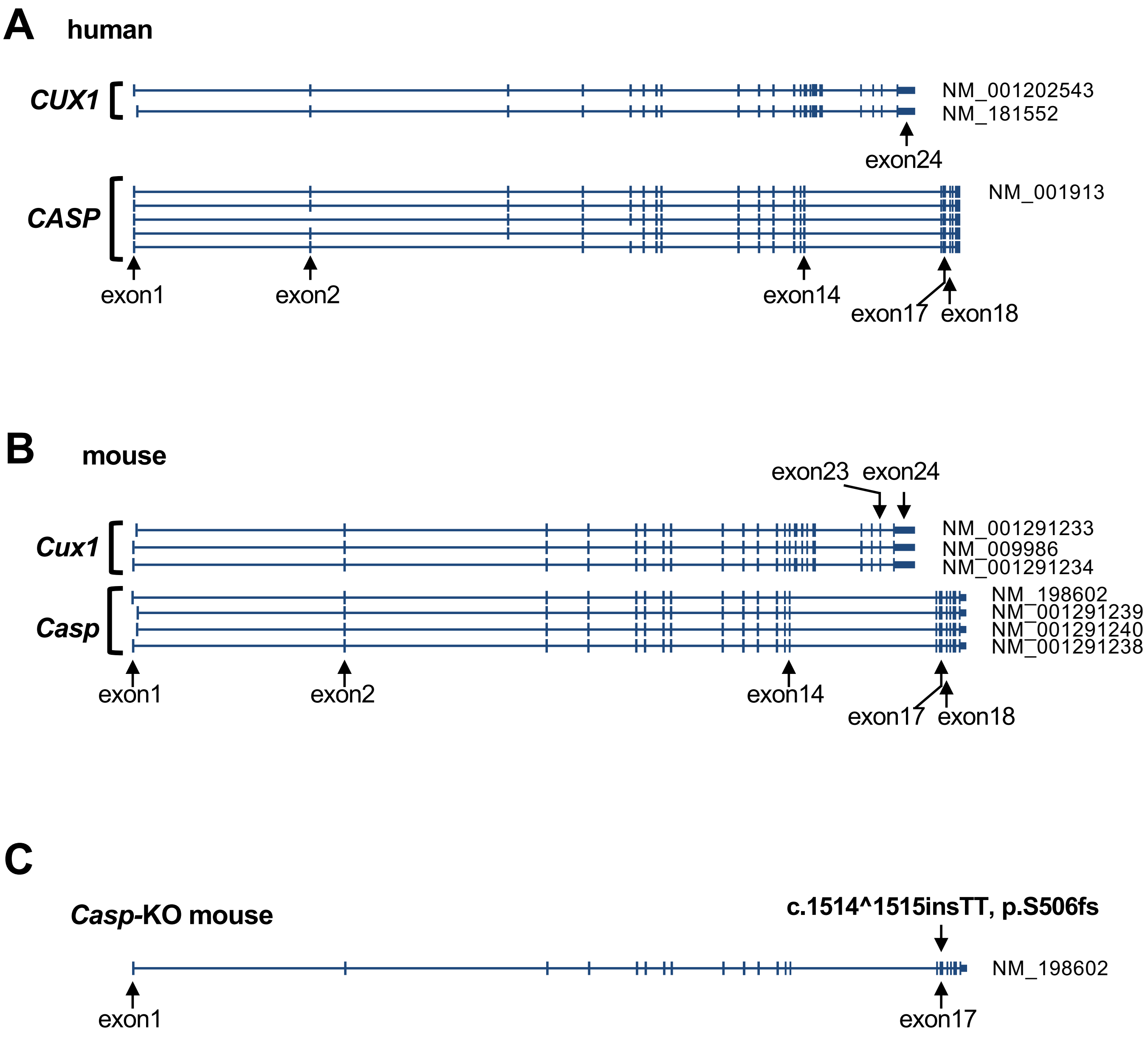


**Figure S7. Genome structures of human and mouse *CUX1* and *CASP* in WT and *Casp*-specific knock-out mice.** Exons 1~14 are commonly found in human *CUX1* and *CASP* in human (**A**) and in the corresponding genes in mice (**B**). In *Casp*-deficient mice, a mutation (c.1514^1515insTT, p.S506fs) was inserted in *Casp*-specific exon 17 (**C**). The diagrams were reconstructed from that of UCSC Genome Browser. Vertical lines indicate exons.

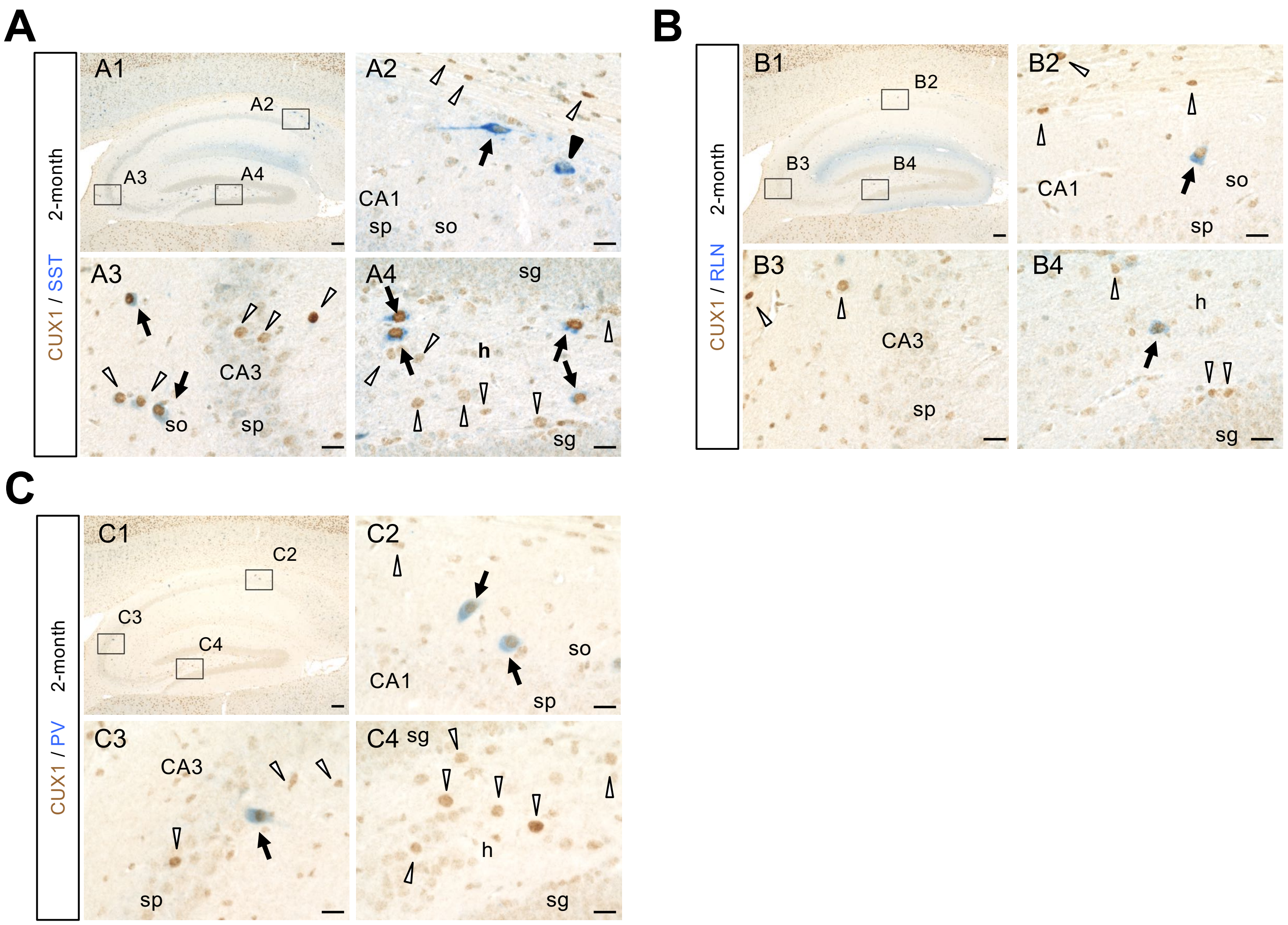


**Figure S8. CUX1 is expressed in hippocampal SST-positive, RLN-positive, or PV-positive interneurons.** CUX1 (brown) is expressed in SST-positive (**A**), RLN-positive (**B**) and PV-positive (**C**) interneurons (arrows). Some of intense CUX1-positive cells were SST-negative, RLN-negative or PV-negative (white arrowheads), and some of SST-positive, RLN-positive or PV-positive cells were CUX1-negative (black arrowhead and not shown). Most of CUX1-positive cells were PV-negative (white arrowhead) in hilus (C). A2-A4, B2-B4, C2-C4: magnified images outlined in A1, B1, C1, respectively. Scale bars = 100 µm (A1, B1 and C1) or 20 µm (A2-A4, B2-B4 and C2-C4). sp; stratum pyramidale, so; stratum oriens, sg; stratum granulosum, h; hilus.

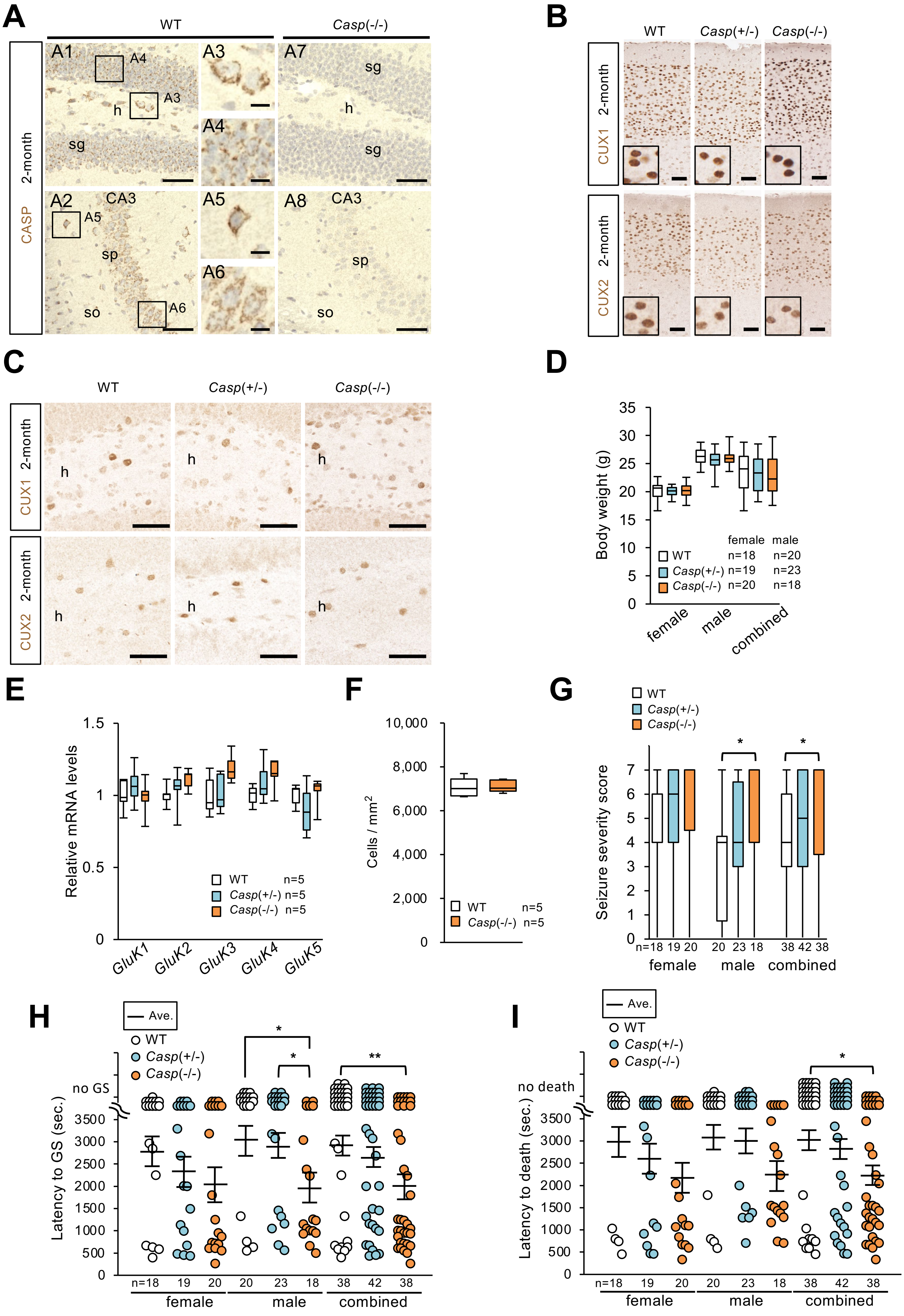

A
WT
Casp(-/-)
2-month
CASP
A1
A2
A3
A4
A5
A6
A7
A8
sg
h
sp
so
CA3
B
WT
Casp(+/-)
Casp(-/-)
CUX1
CUX2
C
D
Body weight (g)
female
male
combined
WT n=18 n=20
Casp(+/-) n=19 n=23
Casp(-/-) n=20 n=18
E
Relative mRNA levels
GluK1
GluK2
GluK3
GluK4
GluK5
WT n=5
Casp(+/-) n=5
Casp(-/-) n=5
F
Cells / mm2
WT n=5
Casp(-/-) n=5
G
Seizure severity score
n=18 19 20 20 23 18 38 42 38
H
Ave.
no GS
Latency to GS (sec.)
I
no death
Latency to death (sec.)

**Figure S9. CASP, CUX1 and CUX2 expression and increased seizure susceptibility to kainate with a tendency of increase in excitatory cell number in entorhinal cortex of *Casp*-deficient mice.** (**A**) CASP immunosignals were observed in both excitatory and inhibitory neurons in the hippocampus, though the signals in inhibitory neurons were especially intense such as those at hilus (A3) and stratum oriens of CA3 (A5), while signals in excitatory neurons are tiny such as those at stratum granulosum (A4) or pyramidal of CA3 (A6). CASP signals well disappeared in *Casp*(-/-) mice (A7, A8). Nuclei were stained with hematoxylin (blue). (**B, C**) Unaltered expression and subcellular localization of CUX1 and CUX2 in cerebral cortex (B) and hilus of hippocampus (C) of *Casp*-deficient mice. (**D**) Body weights were similar among *Casp*-deficient mice and WT littermates at 2-months. (**E**) qPCR experiments showed that mRNA expression levels of kainate receptor subunits were not altered in *Casp*-deficient mice at 2-month-old. (**F**) In a Nissl staining, number of entorhinal cortex layer II-III neurons was comparable between WT and *Casp*(-/-) mice (2-month-old). (**G**) Seizure severity scores were significantly higher in *Casp*(-/-) male and combined gender mice. **(H)** Latencies to onset of generalized seizures (GS) were significantly decreased in male and combined gender of *Casp*(-/-) mice. (**I**) Latencies until death were also significantly decreased in combined gender of *Casp*(-/-) mice. Mice without GS or death within 3,600 sec were plotted at "no GS" or "no death", respectively. Circles represent individual mice. One-way ANOVA followed by Tukey–Kramer Multiple Comparison Test (D, E, G-I), or one-way ANOVA test (F). mean (horizontal bars) ± s.e.m. (H, I). n=mouse number. Scale bars= 50 µm (A1, A2, A7, A8, B and C) or 10 µm (A3-A6). * $P < 0.05$, ** $P < 0.01$.

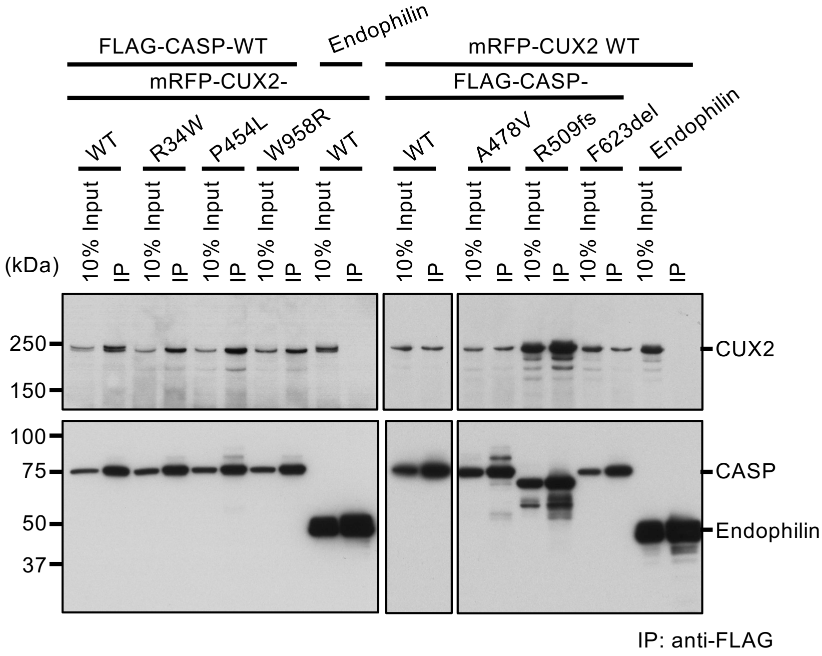


**Figure S10. CASP interacts with CUX2.** mRFP-tagged CUX2 was co-immunoprecipitated with FLAG-tagged CASP. Mutations did not affect the binding. Endophilin: negative control. Original blots are presented in Supplementary Figure S11.

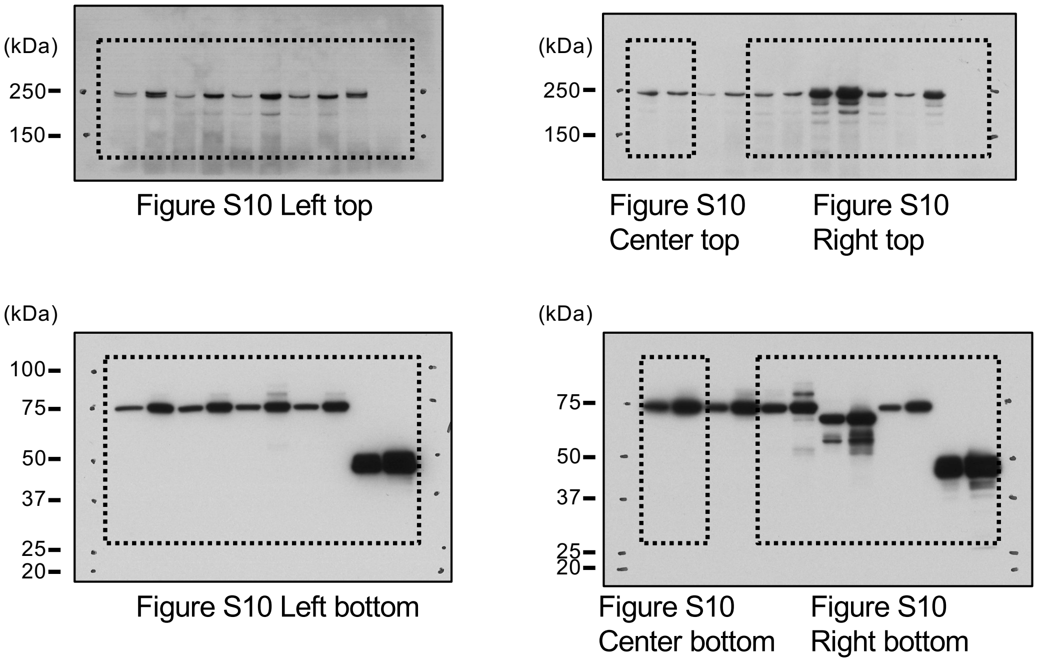


**Figure S11. Full size western blot images.** Original scanned western blot data that were used to generate Figure S10. Dashed rectangles in the images indicate the location of the cropped images. Images of blots with adequate length and membrane edges could not be provided because the blots were cut prior to hybridisation with antibodies and scanned at inside of blots, respectively.
